# Quasi-periodic patterns of brain activity in individuals with Attention-Deficit/Hyperactivity Disorder

**DOI:** 10.1101/435347

**Authors:** Anzar Abbas, Yasmine Bassil, Shella Keilholz

**Affiliations:** Neuroscience, Emory University, 1760 Haygood Dr NE W200, Atlanta, GA 30322; College of Sciences, Georgia Institute of Technology, 225 North Ave, Atlanta, GA 30332; Biomedical Engineering, Emory University and Georgia Institute of Technology, 1760 Haygood Dr NE W200, Atlanta, GA 30322

**Keywords:** attention-deficit/hyperactivity disorder, resting-state fMRI, functional connectivity, default mode network, task positive network, quasi-periodic patterns, dynamic functional connectivity

## Abstract

Individuals with attention-deficit/hyperactivity disorder have been shown to have disrupted functional connectivity in the default mode and task positive networks. Traditional fMRI analysis techniques that focus on ‘static’ changes in functional connectivity have been successful in identifying differences between healthy controls and individuals with ADHD. However, such analyses are unable to explain the mechanisms behind the functional connectivity differences observed. Here, we study dynamic changes in functional connectivity in individuals with ADHD through investigation of quasi-periodic patterns (QPPs). QPPs are reliably recurring low-frequency spatiotemporal patterns in the brain linked to infra-slow electrical activity. They have been shown to contribute to functional connectivity observed through static analysis techniques. We find that QPPs contribute to functional connectivity specifically in regions that are disrupted during ADHD. Individuals with ADHD also show differences in the spatiotemporal pattern observed within the QPPs. This difference results in a weaker contribution of QPPs to functional connectivity in the default mode and task positive networks. We conclude that quasi-periodic patterns provide insight into the mechanisms behind functional connectivity differences seen in individuals with ADHD. This allows for a better understanding of the etiology of the disorder and development of effective treatments.

**Highlights:**

- Default mode and task positive network connectivity is disrupted in ADHD
- Quasi-periodic patterns contribute to typical functional connectivity in the DMN and TPN
- The contribution of QPPs to functional connectivity is diminished in individuals with ADHD
- QPPs could explain BOLD dynamics underlying static functional connectivity differences in ADHD

## 1 Introduction

Attention-deficit/hyperactivity disorder (ADHD) is the most commonly diagnosed neurodevelopmental disorder among children and adolescents in the United States (Subcommittee on Attention-Deficit/Hyperactivity Disorder, 2011). Changing attitudes towards the diagnosis of ADHD are leading to a further increase in its prevalence worldwide (Davidovich et al., 2017). ADHD is characterized by pervasive levels of inattention, hyperactivity, and impulsivity (American Psychiatric Association, 2013). It can lead to difficulties in personal and academic endeavors (Bagwell et al., 2001; Barkley et al., 1991) and cause significant burden on families and society (Matza et al., 2005). Understanding the pathophysiology behind ADHD is crucial for the development of effective treatments.

Etiological models of brain disorders such as ADHD are shifting from focusing on individual brain regions to prioritizing the investigation of large-scale network interactions across the brain (Raj et al., 2018; Konrad & Eickhoff, 2010). As a consequence, non-invasive whole-brain imaging methods are playing an important role in understanding the etiology of brain disorders (Wintermark et al., 2018; Weyandt et al. 2013). Notably, functional magnetic resonance imaging (fMRI) and the blood oxygenation level dependent (BOLD) signal has been critical in studying network interactions in the brain and how they can be disrupted (Stam et al., 2014). Correlation of BOLD signal from anatomically distinct brain regions over time is assumed to be an indication of functional connectivity between those regions. Individuals with brain disorders, such as ADHD, often show altered functional connectivity in the brain (Konrad & Eickhoff et al., 2010; Cortese et al., 2012; Hart et al., 2012).

Such disruptions in functional connectivity have been a central focus of a number of studies on brain disorders (for review, see Du et al., 2018), including ADHD (Konrad & Eickhoff, 2010). Findings from these studies have assisted in identifying brain regions and functional networks relevant to understanding the etiology of brain disorders. However, most of these studies have relied on traditional analyses of functional connectivity, which assume a stationary relationship between brain regions over the course of an fMRI scan (5-10 minutes or longer). Such ‘static’ analyses of functional connectivity fail to consider the rich time-varying component of the BOLD signal present in the data (Chang et al., 2010; Du et al., 2018). Hence, more recent fMRI studies have focused on dynamic analysis of the BOLD signal to better understand network interactions over time. This can help uncover the cause of functional connectivity disruptions seen in individuals with brain disorders (Hutchinson et al., 2013).

ADHD is associated with dysfunction in the default mode network of the brain (DMN) (Castellanos et al., 2008; Uddin et al., 2008) and its relationship with the task positive network (TPN) (Tian et al., 2006; Wang et al., 2008; Wolf et al., 2009; Rubia et al., 2009a; for reviews on structural and functional connectivity disruptions in individuals with ADHD, see Konrad & Eickhoff, 2010; Cortese et al., 2012; Hart et al., 2012). Though there remains uncertainty on the directionality of some observed differences, evidence has predominantly converged on the relevance of the DMN and TPN when studying functional connectivity in individuals with ADHD. This also aligns with the prevailing understanding that DMN-TPN interactions are relevant for attentional control and vigilance (Fox et al., 2005; Raichle, 2015; Thompson et al., 2013). An investigation of the dynamics of these functional networks may help further the understanding of functional connectivity differences seen in individuals with ADHD.

Our group has reported a reliably observable low-frequency spatiotemporal pattern in the brain that involves the DMN and TPN (Majeed et al., 2009; Majeed et al., 2011; Thompson et al., 2014; Belloy et al., 2018; Yousefi et al., 2018; Abbas et al., 2018a). The pattern lasts approximately 20 seconds in humans. It involves an initial increase in BOLD signal in the DMN accompanied by a decrease in BOLD signal in the TPN. This is followed by a subsequent decrease in BOLD signal in the DMN alongside an increase in BOLD signal in the TPN. The sequence of events, which capture the strong anti-correlation between the DMN and TPN (Fox et al., 2005), occurs quasi-periodically over the course of a functional scan. Hence, it has been referred to as a quasi-periodic pattern (QPP). QPPs have been observed in mice (Belloy et al., 2018), rats (Majeed et al., 2009; Majeed et al, 2011; Thompson et al., 2014), rhesus macaques (Abbas et al., 2016), and in resting-state and task-performing humans (Majeed et al., 2011; Abbas et al., 2018a). QPPs are correlated with infra-slow neural activity (Thompson et al., 2014) and are distinct from physiological noise and global signal (Yousefi et al., 2018). Most importantly for the scope of this study, QPPs have been shown to contribute to functional connectivity in the DMN and TPN (Abbas et al., 2018a). This makes them highly relevant to functional connectivity differences seen in individuals with ADHD.

It may be that QPPs are contributing to functional connectivity differences in the connections typically disrupted during ADHD. Such a conclusion would further the understanding of the dynamic processes involved in the etiology of the disorder. In this study, we first create masks of the DMN and TPN in healthy controls and adolescents with ADHD. Next, we search for functional connectivity differences between the Control and ADHD groups. We then apply a pattern-finding algorithm to search for QPPs in both groups, and differentiate between the spatiotemporal patterns that are observed. Finally, we use regression to remove the QPPs from the functional scans in each group and measure their contribution to functional connectivity in the DMN and TPN. Our findings confirm functional connectivity differences previously observed in individuals with ADHD. Notably, we show that QPPs contribute to functional connectivity in the brain in regions relevant to ADHD. This is the first investigation of QPPs in any individuals with a brain disorder and suggest a role of QPPs in maintaining a healthy functional architecture of the brain.

## 2 Methods

The Matlab script used for all analysis and figures included in this study is available on github.com/anzarabbas/qppsadhd.

### 2.1 Data acquisition and preprocessing

All resting-state data was downloaded from the ADHD-200 Sample, accessible through the 1000 Functional Connectomes Project (ADHD-200 Consortium, 2012; Biswal et al. 2010). Within the ADHD-200 Sample, the New York University, Peking University, and NeuroImage datasets were used. These datasets were selected based on the similarity of their scan parameters and availability of diagnostic information and data quality control assessments. An overview of scan acquisition parameters for each dataset is provided in Table 1.

**Table 1.**
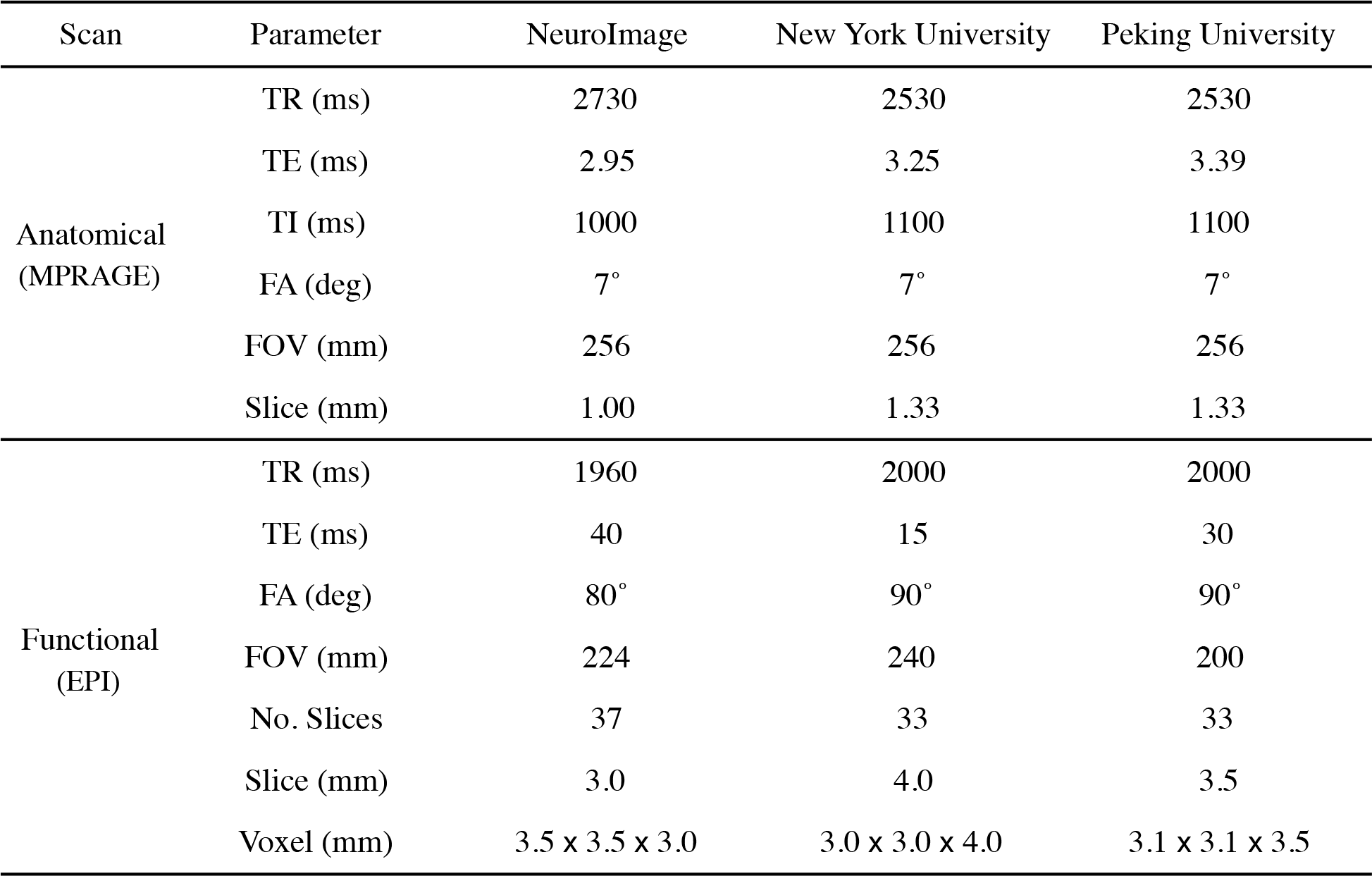
Anatomical and functional scan parameters for the ADHD 200 Sample datasets used in the study. In all cases, the anatomical scans were acquired through a T1-weighted 3D magnetization-prepared rapid gradient echo (MPRAGE) sequence and the functional scans were acquired through a gradient echo-planar imaging (EPI) sequence.

| Scan | Parameter | NeuroImage | New York University | Peking University |
| --- | --- | --- | --- | --- |
| Anatomical<br>(MPRAGE) | TR (ms) | 2730 | 2530 | 2530 |
|  | TE (ms) | 2.95 | 3.25 | 3.39 |
|  | TI (ms) | 1000 | 1100 | 1100 |
|  | FA (deg) | 7° | 7° | 7° |
|  | FOV (mm) | 256 | 256 | 256 |
|  | Slice (mm) | 1.00 | 1.33 | 1.33 |
| Functional<br>(EPI) | TR (ms) | 1960 | 2000 | 2000 |
|  | TE (ms) | 40 | 15 | 30 |
|  | FA (deg) | 80° | 90° | 90° |
|  | FOV (mm) | 224 | 240 | 200 |
|  | No. Slices | 37 | 33 | 33 |
|  | Slice (mm) | 3.0 | 4.0 | 3.5 |
|  | Voxel (mm) | 3.5 x 3.5 x 3.0 | 3.0 x 3.0 x 4.0 | 3.1 x 3.1 x 3.5 |

Each dataset contained MRI scans from healthy children, adolescents, and some adults and individuals diagnosed with ADHD. Of the three main sub-types of ADHD (Inattentive, Hyperactive-Impulsive, and Combined), only individuals diagnosed with the Combined ADHD sub-type were used in this study. The selection of one sub-type was intended to reduce variability in the results and the Combined sub-type provided the largest dataset among the three. Though the ADHD group had a combination of treated and medically naive individuals, all participants had been removed from any psycho-stimulant medication 24-48 hours prior to collection of functional data. For all individuals, only scans that had passed the ADHD 200 Sample quality control assessment were used. For individuals that had more than one functional scan, only the first scan was used in the study. In the end, the Control group contained 106 healthy individuals (age range 7-26, *μ* = 14.6 years ± 3.8; 56 females) and the ADHD group contained 106 individuals with the Combined sub-type of ADHD (age range 7-21, *μ* = 12.6 ± 3.3; 10 females). Of the 106 individuals in each group, 22 were from the NeuroImage Sample, 57 were from the New York University dataset, and 33 were from the Peking University dataset.

All preprocessing was conducted using FSL 5.0 (Jenkinson et al., 2012) and MATLAB (Mathworks, Natick, MA). Anatomical data was registered to the 2 mm Montreal Neurological Institute (MNI) brain atlas using FLIRT (Jenkinson & Smith, 2001; Jenkinson et al., 2002), skull-stripped using BET, and tissue segmented into white matter, gray matter, and cerebrospinal fluid using FAST (Zhang et al., 2001). Prior to carrying out preprocessing steps on the functional data, all functional scans in both groups were shortened to 150 timepoints so that all scans could be of the same length. Functional data was slice time corrected using FSL’s slicetimer tool, motion corrected using MCFLIRT (Jenkinson et al., 2002), registered to MNI space using FLIRT, and spatially smoothed with a 6 mm Gaussian kernel using FSLMATHS. MATLAB was used to apply a Fast Fourier Transform bandpass temporal filter between 0.01 and 0.08 Hz. Lastly, global, white matter, and cerebrospinal signals were regressed and all voxel timecourses were z-scored. The automated preprocessing pipeline used in this study is available for use by other researchers on github.com/anzarabbas/fmripreprocess.

### 2.2 Acquisition of default mode and task positive networks

A data-driven method was used to acquire masks of the default mode and task positive networks from the Control and ADHD groups. For each group, 30 functional scans were concatenated in time. The 30 scans selected had 10 scans each from the three datasets (NeuroImage, New York University, and Peking University) to ensure results were not biased by any one dataset. The average BOLD signal over time, or the mean timecourse, of the posterior cingulate cortex (PCC) was calculated from the concatenated scans. Pearson correlations were then conducted between the mean timecourse of the PCC and the timecourse of every voxel in the brain. The 10% of voxels that were most correlated with the PCC were labeled as the DMN. The 10% of voxels that were most *anti-correlated* with the PCC were labeled as the TPN (Fox et al., 2005).

The functional scans were segmented into 273 regions of interest (ROIs) from the Brainnetome ROI atlas (Fan et al., 2016). For each ROI, the binary mask of the ROI was multiplied by the binary masks of the DMN and TPN to check for any spatial overlap between the ROI and either network. If the ROI contained voxels that were also part of either the DMN or TPN, the number of such voxels were counted and their mean correlation with the PCC was recorded. By doing so, a list of ROIs in the DMN and TPN was constructed, which contained information on how much the ROI overlapped with the DMN or TPN and how strongly it was correlated or anti-correlated with the PCC (Supplementary Table 1 for the DMN and Supplementary Table 2 for the TPN). The list was used to compare the ROIs included in the DMN and TPN masks acquired from the Control and ADHD groups. It was also used to compare the correlation strength of the ROIs with the PCC across the Control and ADHD groups. Finally, the DMN and TPN masks were used to acquire mean timecourses of the DMN and TPN from every scan in each group. The overall anticorrelation between the mean timecourse of the DMN and the mean timecourse of the TPN across all scans was compared between the Control and ADHD groups using a Mann-Whitney U test.

### 2.3 Acquisition of quasi-periodic patterns

A spatiotemporal pattern-finding algorithm, described in Majeed et al. (2011), was used to search for repeating patterns in the functional scans. The method in which the algorithm was applied and all parameters used are outlined in Abbas et al. (2018).

The pattern-finding algorithm begins by conducting a sliding correlation between a random, user-defined starting segment within a functional timeseries with the functional timeseries itself. If the brain activity captured in the segment repeats at other instances in the functional timeseries, the resulting sliding correlation vector will contain local maxima, or peaks, indicating those occurrences. At each of those instances, additional segments of the same length are extracted and averaged together into an updated segment. Subsequent sliding correlations are then conducted between the continually updated segment and the functional timeseries. These steps are repeated until the updated segment no longer shows variation and represents a reliably repeating pattern of brain activity within the functional timeseries. The algorithm has two main outputs: A repeating spatiotemporal pattern from within the functional timeseries, and a sliding correlation vector of the pattern with the functional timeseries itself.

As mentioned in the Introduction, the quasi-periodic pattern being investigated in this study lasts approximately 20 seconds. It involves an initial increase in BOLD signal in the DMN alongside a decrease in BOLD signal in the TPN, followed by a subsequent decrease in BOLD signal in the DMN alongside an increase in BOLD signal in the TPN. Concisely stated, QPPs involve a propagation of BOLD signal from the DMN to the TPN, or a DMN/TPN switch in BOLD activation (Abbas et al., 2018a; Majeed et al., 2011; Yousefi et al., 2018). The pattern-finding algorithm described above has been shown to output such a pattern. However, the phase in which the DMN/TPN switch occurs within the pattern can vary depending on the user-defined starting segment (Yousefi et al., 2018). To ensure that the DMN/TPN switch occurs in the same phase in the QPPs that will be acquired from both groups, the algorithm is run multiple times using randomly selected starting segments.

For the Control and ADHD groups separately, 30 functional scans were again concatenated (10 scans from each dataset). The pattern-finding algorithm was applied to the concatenated timeseries 100 times with unique, randomly-selected starting segments. The resulting 100 patterns outputted by the algorithm for each group were analyzed for a DMN-to-TPN transition in BOLD activation. The pattern most closely matching a DMN-to-TPN switch was selected and designated as a representative QPP for its respective group. By doing so, one representative QPP was established for the Control group and another representative QPP was established for the ADHD group. It is unlikely that the 30 scans concatenated before application of the algorithm biased the spatiotemporal pattern captured in the QPP. It has been previously shown that 25 concatenated scans (of similar length) were sufficient in removing variability in the pattern outputted by the algorithm (Abbas et al., 2018a). It has also been shown that QPPs acquired from concatenated data are the same as averaged QPPs from individual scans (Yousefi et al., 2018).

The spatiotemporal pattern captured in the QPP was compared between the Control and ADHD groups. The QPPs were segmented into the 273 ROIs in the Brainnetome ROI atlas. The mean timecourse of each ROI was calculated for both QPPs. For each ROI, a Pearson correlation was conducted between its mean timecourse in the Control QPP and its mean timecourse in the ADHD QPP. The resulting values for all 273 ROIs were compiled into Supplementary Table 3. Strong correlation of an ROI’s timecourse in the two QPPs indicates that the ROI behaved similarly in both groups’ QPPs. Anti-correlation of an ROI’s timecourse in the two QPPs indicates that the ROI behaved differently in the QPP acquired from individuals with ADHD.

Next, the strength and frequency of the QPPs was compared between groups. Sliding correlations of the Control and ADHD QPPs were conducted with all functional scans in their respective groups. The resulting sliding correlation vectors contained local maxima, or peaks, in correlation, which signified the occurrence of QPPs at those instances in the functional scans. The strength of the QPP was defined as the mean height of those peaks. The frequency of the QPP was defined as the rate of occurrence of those peaks over time. In this study, frequency was measured in peaks per minute. To compare the strength and frequency of the QPPs across groups, an arbitrary peak height threshold of 0.1 was chosen. First, the mean height of all peaks greater than the threshold was compared between the Control and ADHD groups. Second, the overall frequency of all peaks greater than the threshold was compared between the Control and ADHD groups. Finally, the arbitrary 0.1 threshold was discarded and the cumulative sliding correlations of the Control and ADHD QPPs with their respective functional scans were plotted as histograms. The distribution of values observed in these histograms were compared between the Control and ADHD groups using a Kolmogorov-Smirnov test.

### 2.4 Removal of QPPs from functional scans

To study the contributions of the Control and ADHD QPPs to functional connectivity in the DMN and TPN, they were removed from the BOLD signal using the regression method described in Abbas et al. (2018). The native QPP from all functional scans from each group was regressed from the BOLD signal. Native QPPs are defined as the QPPs acquired from that group. For example, the Control QPP is native to all the functional scans in the Control group. For each functional scan, a unique regressor was calculated for every brain voxel: The sliding correlation of the QPP was convolved with the timecourse of each brain voxel during the QPP. The obtained regressor was z-scored to match the signal in the functional scan. Next, linear regression was carried out using standardized/beta coefficients and the regressors calculated for each brain voxel. By doing so, a functional scan with attenuated presence of the QPP in the BOLD signal was produced. The efficacy of this regression method was demonstrated by conducting subsequent sliding correlations of the QPPs with all QPP-regressed functional scans. The same comparison of strength and frequency of QPPs described in the last paragraph of Section 2.3 was conducted in the QPP-removed functional scans. Differences in the strength and frequency of QPPs after their removal were compared.

### 2.5 Analysis of functional connectivity

Before analysis of functional connectivity, a new set of ROIs focused only on regions in the DMN and TPN were created. First, the 273 ROIs from the Brainnetome atlas were consolidated into 26 ROIs based on the structural hierarchy of the atlas. For example, the 14 ROIs within the superior frontal gyrus were consolidated into a single ROI depicting the entire superior frontal gyrus. Next, the binary mask of each consolidated ROI was multiplied by the binary masks of the DMN and TPN to check for any spatial overlap between the consolidated ROI and either network. Only ROIs that had spatial overlap with either network were included in the new atlas. Within these ROIs, only the voxels that overlapped with the DMN or TPN were included. For example, the superior frontal gyrus has a total of 11341 voxels. Of these, 3093 voxels overlapped with the DMN mask. Only those 3093 voxels were included in the DMN’s superior frontal gyrus ROI. However, 1253 separate voxels in the superior frontal gyrus overlapped with the TPN mask. Those 1253 voxels were included in the TPN’s superior frontal gyrus ROI. In the end, the new set of ROIs contained a total of 36 ROIs, half of which were DMN ROIs and half of which were TPN ROIs. Since there were differences in the DMN and TPN masks acquired from the Control and ADHD groups, a union of the DMN and TPN masks from the two groups was used during the construction of the new set of ROIs.

Functional connectivity matrices were created to visualize the strength of connectivity within and across the DMN and TPN in both groups. For each functional scan, one functional connectivity matrix was created before QPP regression, and one functional connectivity matrix was created after its native QPP had been regressed. To create each of these matrices, the Pearson correlation between the mean timecourse of each ROI in the functional scan and the mean timecourse of all other ROIs in the functional scan was calculated. The values from each of the correlations were Fischer z-transformed and arranged into a 36 ROI × 36 ROI matrix. The functional connectivity matrices from all scans were averaged to obtain the mean functional connectivity for that group. In the end, each group had a mean functional connectivity matrix both before and after regression of its native QPP.

The newly created functional connectivity matrices were used to compare the strength of functional connectivity between ROIs in the DMN and TPN. First, functional connectivity strength was compared between the Control and ADHD groups. This was done once before regression of any QPPs, and again after regression of native QPPs from the functional scans. The functional connectivity differences observed between the two groups before and after regression of native QPPs were compared. Second, functional connectivity strength was compared within the Control and ADHD groups after removal of their native QPP. The effect of the regression of the native QPPs on functional connectivity strength was then compared between the Control and ADHD groups. To conduct all comparisons, a two-sample *t*-test was performed for each ROI connection to check for a significant change in functional connectivity strength. Given that there were 648 connections to compare, multiple comparisons correction was performed using the false detection rate correction method presented in Benjamin and Hochberg (1995). For all connections that were significantly different in functional connectivity strength, the mean difference in connectivity was displayed in the same style as the functional connectivity matrices.

## 3 Results

### 3.1 Differences in DMN and TPN masks between groups

Masks of the default mode and task positive networks acquired from the Control and ADHD groups were largely similar (Figure 1a; Figure 1b). A full list of ROIs in the Brainnetome atlas that overlapped with either the DMN or TPN is shown in Supplementary Tables 1 and 2. The tables also list the number of voxels in each ROI that overlapped with the DMN or TPN, and the mean correlation strength between the overlapping voxels in each ROI and the PCC.

**Figure 1.**
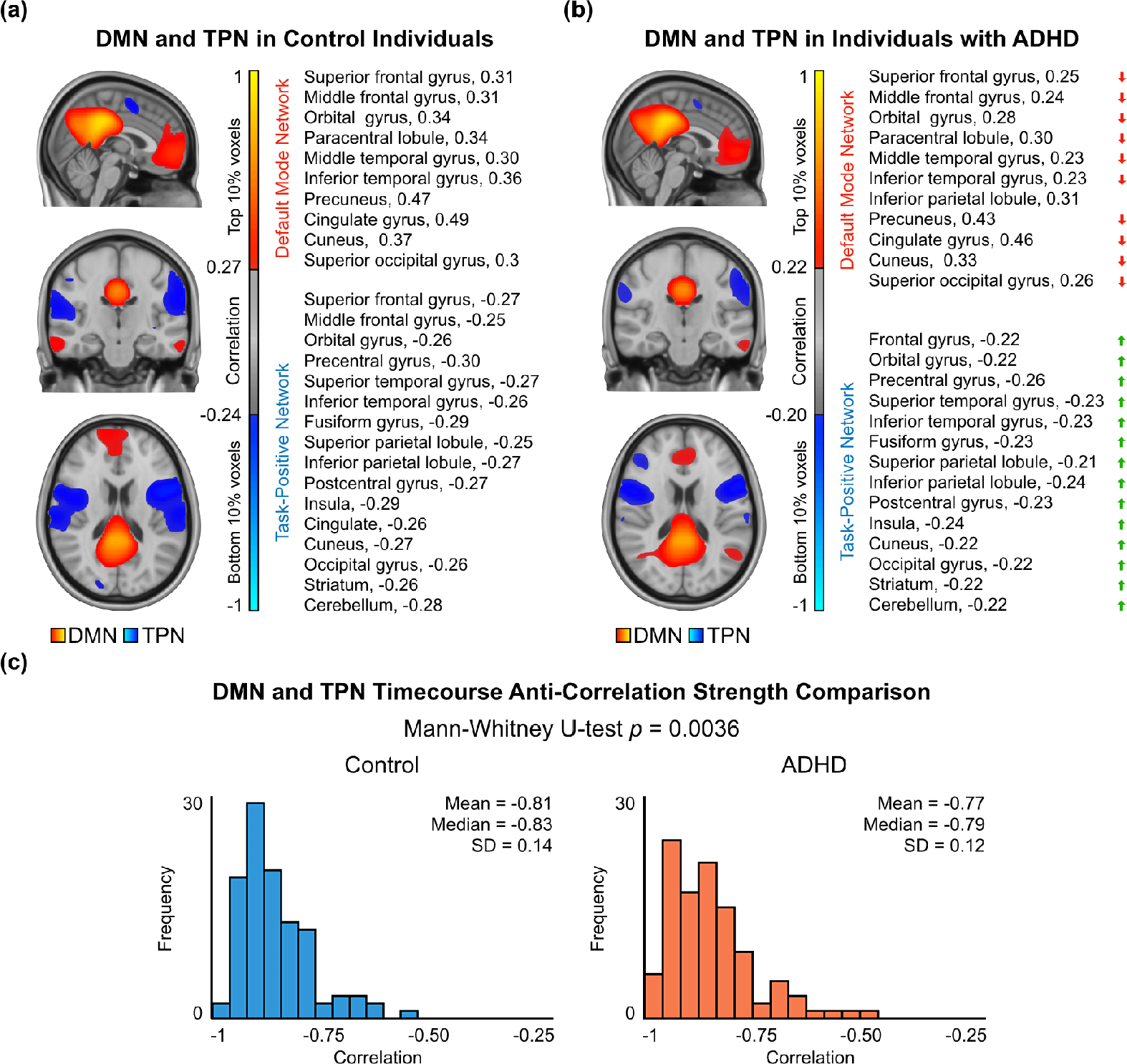
DMN and TPN in the Control and ADHD groups. Correlation between the mean timecourse of the PCC and every voxel in the brain was calculated. The 10% of voxels most and least correlated with the PCC were defined as the DMN and TPN respectively. **(a)** *Left*: The DMN and TPN in the Control group. The DMN comprises all voxels that had correlation with the PCC > 0.27. The TPN comprises all voxels that had correlation with the PCC < −0.24. *Right*: Names of regions in the DMN and TPN in the Control group. **(b)** *Left*: The DMN and TPN in the ADHD group. The DMN comprises all voxels that had correlation with the PCC > 0.22. The TPN comprises all voxels that had correlation with the PCC < −0.20. *Right*: Names of regions in the DMN and TPN in the ADHD group. A full list of ROIs in the DMN and TPN, including subdivisions, number of voxels, and strength of correlation with PCC is provided in Supplementary Tables 1 and 2. Compared to the Control group, areas in the DMN had overall lower correlation with the PCC, while areas in the TPN had overall weaker anti-correlation with the PCC. **(c)** Distributions of anti-correlation strength between DMN and TPN timecourses in all Control (*left*) and ADHD (*right*) scans. Given the non-parametric distributions, a Mann-Whitney U-test was performed to compare the strength of anti-correlation, which showed weaker anti-correlation in the ADHD group compared to the control group (*p* = 0.0036).

For both the Control and ADHD groups, the DMN included regions in the superior and middle frontal gyri, orbital gyrus, paracentral lobule, middle and inferior temporal gyri, inferior parietal lobule, precuneus, cingulate, cuneus, superior occipital gyrus, hippocampus, and cerebellum. In the ADHD group, the DMN also included regions in the superior temporal gyrus, superior parietal lobule, striatum, and thalamus. Though DMN ROIs unique to the ADHD group are considered in all analyses, the number of voxels in those ROIs that overlapped with the DMN mask was relatively low (< 10 voxels for each ROI, as opposed to a mean of 1461 ± 1464 voxels for the other ROIs in the ADHD DMN), with possible exception of the thalamus (33 voxels in the ADHD DMN as opposed to 0 voxels in the Control DMN). Additionally, though the hippocampus and part of the cerebellum were in the DMN mask from both groups, the number of voxels in those ROIs that overlapped with the DMN were also relatively low (< 10 voxels).

For both the Control and ADHD groups, the TPN included regions in the superior, middle, and inferior frontal gyri, orbital gyrus, precentral gyrus, superior temporal gyrus, inferior temporal gyrus, fusiform gyrus, superior and inferior parietal lobules, postcentral gyrus, insula, cuneus, occipital gyrus, striatum, and cerebellum. In the Control group, the TPN also included the cingulate. In the ADHD group, the TPN also included the middle temporal gyrus. Though TPN ROIs unique to either the Control or ADHD groups are considered in all analyses, the number of voxels in those ROIs that overlapped with the TPN mask was relatively low (< 25 for each ROI, as opposed to a mean of 1238 ± 750 voxels and 1163 ± 787 voxels for the other ROIs in the Control and ADHD TPNs respectively). Additionally, the middle frontal and middle temporal gyri had far greater overlap with the ADHD TPN mask (361 and 393 voxels respectively) than they did with the Control TPN mask (24 voxels each).

As described in the Methods, ROIs in the DMN and TPN masks were selected based off their correlation with the PCC. The strength of correlation of the DMN ROIs and the strength of anticorrelation of the TPN ROIs with the PCC was compared between the Control and ADHD groups. For ROIs in the DMN, the mean correlation with the PCC was greater in the Control group (< = 0.37 ± 0.11) than the ADHD group (*μ* = 0.31 ± 0.12; *p* = 0.0066). For ROIs in the TPN, the mean anti-correlation with the PCC was greater in the Control group (*μ* = −0.28 ± 0.02) than the ADHD group (*μ* = −0.23 ± 0.02; *p* = 4.6995e-33).

The strength of anti-correlation between the DMN and TPN timecourses for all scans were also compared between the Control and ADHD groups (Figure 1c). Given anti-correlation values had a skewed distribution, a Mann-Whitney U-test was used to compare the strength of DMN and TPN anti-correlation across groups. The ADHD group had significantly weaker anti-correlation between the DMN and TPN (*μ* = −0.77 ± 0.12, median = −0.79) compared to the Control group (*μ* = −0.81 ± 0.13, median = −0.83; *p* = 0.0036).

### 3.2 Differences in QPPs between groups

Application of the pattern-finding algorithm resulted in the observation of a quasi-periodic pattern lasting approximately 20 seconds in both the Control and ADHD groups (Figure 2a; Figure 2b). The spatiotemporal pattern involved an initial increase in BOLD signal in the DMN accompanied by a decrease in BOLD signal in the DMN. This was followed by a decrease in BOLD signal in the DMN accompanied by an increase in BOLD signal in the TPN. The spatiotemporal pattern and its strength and frequency in functional scans was compared between the Control and ADHD groups.

**Figure 2.**
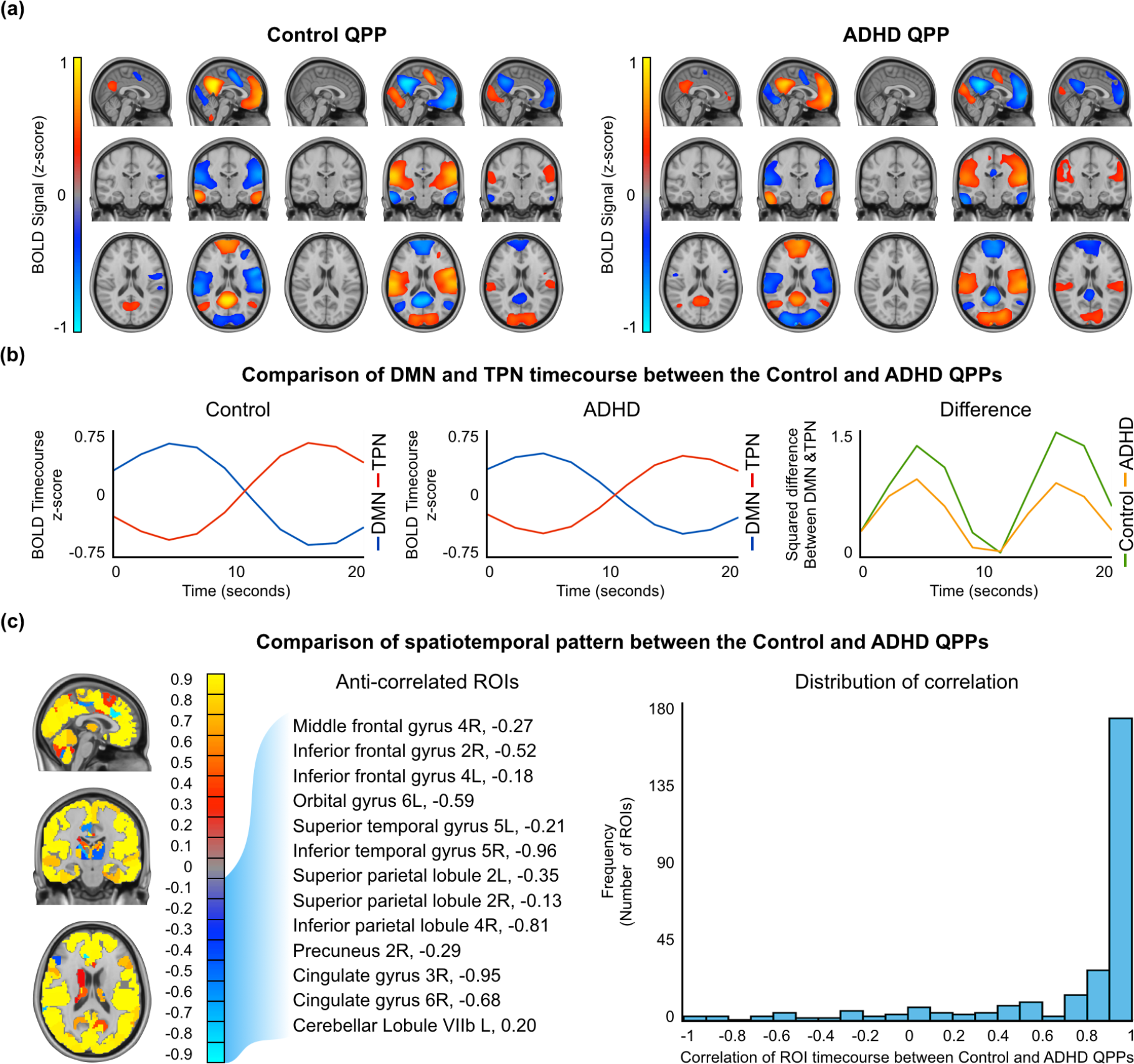
Spatiotemporal comparison of the Control and ADHD QPPs. **(a)** Areas with large increases or decrease in the BOLD signal during the Control (left) and ADHD (*right*) QPPs. Only top and bottom 10% values are shown. **(b)** Timecourse of the DMN and TPN during the Control (*left*) and ADHD (*right*) QPPs. *Right*: The square of the difference between the Control and DMN timecourse at each timepoint in the Control and ADHD QPPs. **(c)** Map of similarities and differences between the Control and ADHD QPPs. Areas of positive correlation are shown in red/yellow. Areas of negative correlation are shown in blue/turquoise. All anti-correlated regions that were also in the DMN and TPN masks in either the Control or ADHD groups are listed. *Right*: Distribution of correlation values for all 273 ROIs shows that most ROI timecourses had > 0.9 correlation between the the two QPPs.

#### 3.2.1 Differences in the spatiotemporal pattern

For each of the 273 ROIs in the Brainnetome ROI atlas, a correlation was performed between the timecourse of the ROI in the Control QPP and its timecourse in the ADHD QPP. The results of all 273 correlations are listed in Supplementary Table 3 and displayed using a colormap in Figure 2c on the left. Overall, the spatiotemporal pattern captured in both QPPs was similar. The distribution of correlation values, shown on the right in Figure 2c, demonstrates that most ROI timecourses were strongly correlated between the Control and ADHD QPPs. A few ROIs had anti-correlated timecourses between the two QPPs. Among them, the ROIs that overlapped with either the DMN or TPN are listed in Table 2 and further explored in the discussion.

**Table 2.** List of regions of interest in the default mode and task positive networks which showed anti-correlated timecourses when comparing quasi-periodic patterns from the Control and ADHD groups. Blue tick marks indicate the overlap of the ROI with the DMN or TPN from the Control group. Red tick marks indicate the overlap of the ROI with the DMN or TPN from the ADHD group. The correlation column shows the strength of anticorrelation between the timecourse of the ROI in the Control and ADHD QPPs.

| Region | DMN | TPN | Correlation |
| --- | --- | --- | --- |
| Middle frontal gyrus, part 4 (ventral area 9/46 ), right |  | ✓ | -0.27 |
| Inferior frontal gyrus, part 2 (inferior frontal sulcus), right |  | ✓ | -0.52 |
| Inferior frontal gyrus, part 4 (rostral area 45), left | ✓ | ✓ | -0.18 |
| Orbital gyrus, part 6 (lateral area 12/47), left | ✓ | ✓ | -0.59 |
| Superior temporal gyrus, part 5 (lateral area 38), left | ✓ | ✓ | -0.21 |
| Inferior temporal gyrus, part 5 (ventrolateral area 37), right | ✓ | ✓ | -0.96 |
| Superior parietal lobule, part 2 (caudal area 7), left |  | ✓ | -0.35 |
| Superior parietal lobule, part 2 (caudal area 7), right |  | ✓ | -0.13 |
| Inferior parietal lobule, part 4 (caudal area 40), right |  | ✓ | -0.81 |
| Precuneus, part 2 (medial area 5), right | ✓ | ✓ | -0.29 |
| Cingulate gyrus, part 3 (pregenual area 32), right | ✓ | ✓ | -0.95 |
| Cingulate gyrus, part 6 (caudal area 24), right | ✓ | ✓ | -0.68 |
| Cerebellar lobule VIIb, left |  | ✓ | -0.20 |

#### 3.2.2 Difference in the DMN and TPN timecourses

Both groups’ QPPs clearly showed a DMN/TPN switch in the spatiotemporal pattern. However, calculating the square of the difference between the DMN and TPN timecourses in each of the QPPs revealed a clear difference in the magnitude of that difference (Figure 2b, *right).* At the two points where DMN and TPN signal was most separated, the mean square difference was 1.4 in the Control group and 0.9 for the ADHD group.

#### 3.2.3 Differences in the strength and frequency

Sliding correlations of the Control and ADHD QPPs were conducted with all functional scans in their respective groups. Examples of the sliding correlation vectors are shown in Figure 3a. The strength of a QPP in a functional scan is defined by the height of the peaks in the sliding correlation vectors. The frequency of a QPP in a functional scan is defined by how often the peaks occur over time. For the purposes of this study, the frequency of a QPP is measured in peaks per minute.

**Figure 3.**
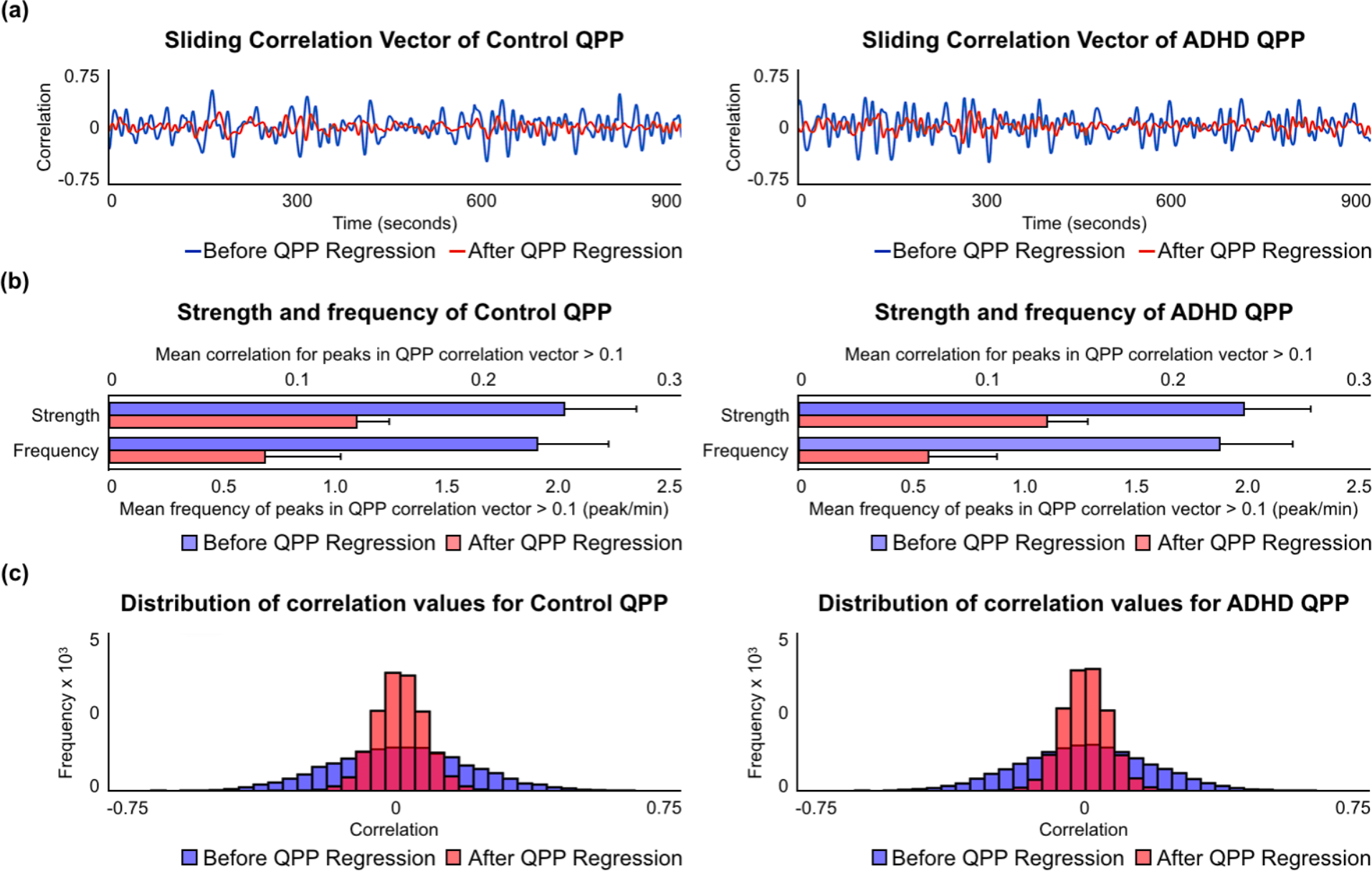
Comparison of the strength and frequency of QPP between the Control and ADHD groups before and after QPP regression. **(a)** Example of sliding correlation vector acquired through sliding correlation of the Control (*left*) and ADHD (*right*) QPPs with three (randomly selected) concatenated functional scans from their respective groups before (*blue*) and after (*red*) native QPP regression **(b)** Strength and frequency of of the Control (*left*) and ADHD (*right*) QPPs compared by setting an arbitrary 0.1 correlation threshold for identifying peaks in the correlation vectors. Top axis shows the strength in correlation and bottom axis shows frequency in peaks per minute before (*blue*) and after (*red*) native QPP regression. **(c)** Strength and frequency of the Control (*left*) and ADHD (*right*) QPPs compared by representing all correlation values in a histogram before (*blue*) and after (*red*) native QPP regression.

For all peaks > 0.1 in correlation, the strength and frequency of the Control QPP in its cumulative sliding correlation with the Control scans (strength *μ* = 0.24 ± 0.04; frequency *μ* = 1.87 ± 0.31 peaks/minute) was similar to the ADHD QPP in its cumulative sliding correlation with the ADHD scans (strength *μ* = 0.23 ± 0.04; frequency *μ* = 1.84 ± 0.32 peaks/minute). The cumulative sliding correlation vectors of the QPPs with each group was also compared without the use of an arbitrary 0.1 threshold by plotting them as histograms (Figure 3e; Figure 3f). Wide, short histograms indicate higher strength and frequency of the QPP, while narrow, tall histograms indicate lower strength and frequency of the QPP. Kolmogorov-Smirnov (KS) tests confirmed that the strength and frequency of the Control and ADHD QPPs in their native scans did not differ. Comparison of the strength of frequency of the QPPs in their non-native scans also did not show any differences (Supplementary Figure 1).

### 3.3 Functional connectivity differences

A functional connectivity matrix displays the strength of functional connectivity in all 648 connections between the 36 brain regions compared in one image, which represents the functional architecture of the DMN and TPN. The matrix has been arranged so that data points closer to the central diagonal show functional connectivity in local connections. Data points further from the central diagonal show functional connectivity in long-range connections. The top-left and bottom-right quadrants show the local functional connectivity in the DMN and TPN respectively. The strength of these connections was expected to be positive as they are depicting functional networks. Alternatively, the top-right and bottom-left quadrants show the functional connectivity between the DMN and TPN. The strength of these connections was expected to be negative, given they are depicting connectivity between anti-correlated networks.

Figure 4a shows the mean functional connectivity in the DMN and TPN in the Control (bottom-left) and ADHD (top-right) groups. Individuals with ADHD showed weaker overall connectivity with the DMN and TPN and weaker anti-correlation across the DMN and TPN. Functional connectivity within DMN ROIs in the ADHD group (*μ* = 0.23 ± 0.40) was weaker than the Control group (*μ* = 0.25 ± 0.45; *p* = 3.72e-39). Functional connectivity within TPN ROIs in the ADHD group (*μ* = 0.26 ± 0.45) was weaker than the Control group (*μ* = 0.31 ± 0.42; *p* = 6.41e-53). Anti-correlation between the DMN and TPN was also weaker in individuals with ADHD (*μ* = −0.21 ± 0.32) compared to the Control group (*μ* = −0.26 ± 0.30; *p* = 3.80e-73).

**Figure 4.**
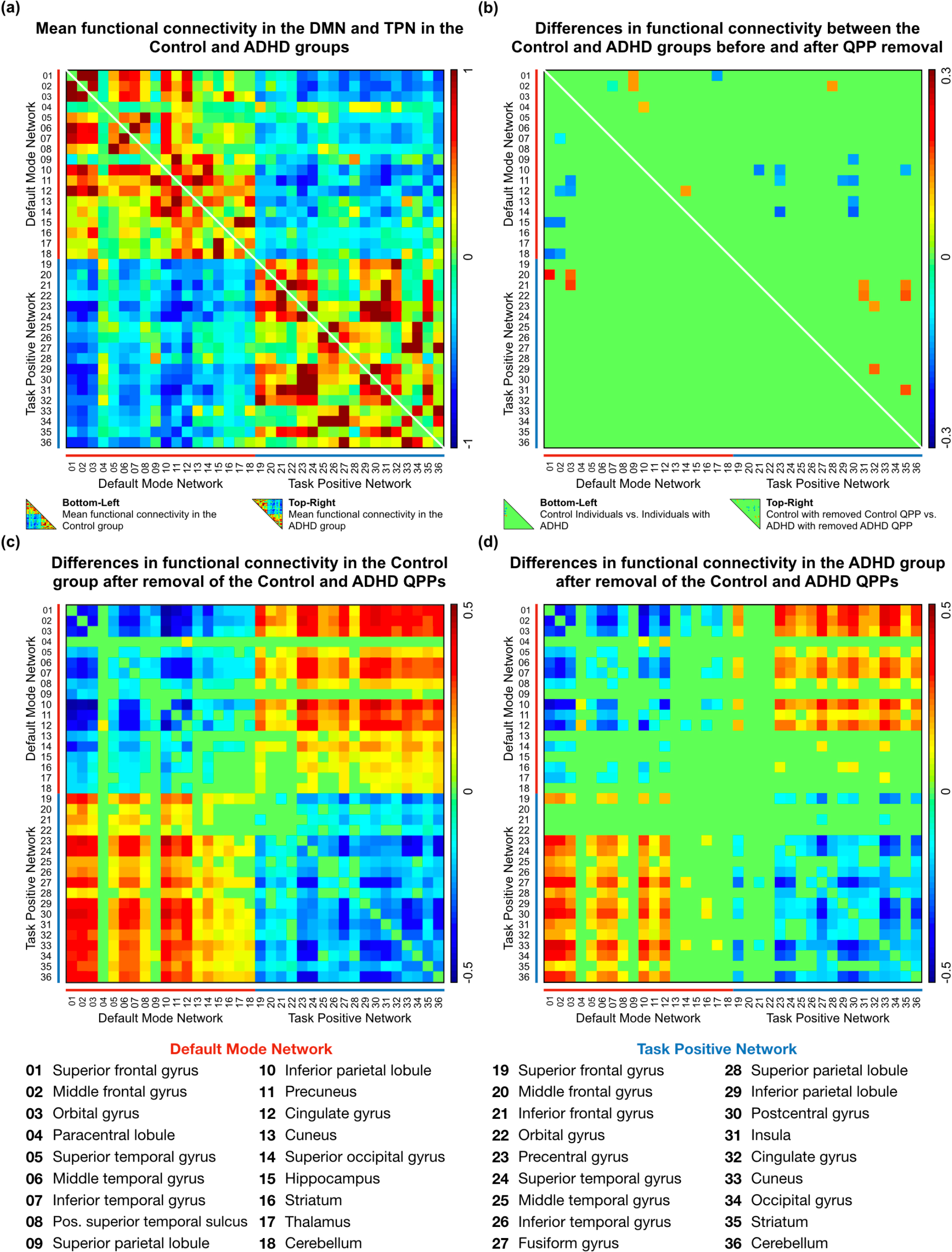
Functional connectivity in 36 ROIs within the DMN and TPN. **(a)** *Bottom-left*: Mean functional connectivity in the Control group. *Top-right*: Mean functional connectivity in the ADHD group. **(b)** *Bottom-left*: Significant differences in functional connectivity between the Control and ADHD groups (*n* = 11). *Top-right*: Significant differences in functional connectivity between the Control and ADHD group after regression of their native QPPs (*n* = 24). **(c)** Significant differences in functional connectivity in the Control group after removal of its native QPP (*n* = 494). **(d)** Significant differences in functional connectivity in the ADHD group after removal of its native QPP (*n* = 280).

Figure 4b shows significant differences in functional connectivity between the Control and ADHD groups. The bottom-left part of the matrix shows differences in functional connectivity before the QPPs were regressed from the functional scans (*n* = 11). Individuals with ADHD showed decreased local functional connectivity in the DMN and decreased anti-correlation between DMN and TPN ROIs. The top-right part of the matrix shows differences in functional connectivity after native QPPs had been regressed from both groups (*n* = 24). The differences were more widespread in this case, but largely comprised on increases in local functional connectivity in the DMN and TPN and increased anti-correlation between the DMN and TPN in individuals with ADHD. These differences are further explored in the discussion.

Figures 4c and 4d show significant differences in functional connectivity in the Control and ADHD groups after regression of their native QPPs. In both groups, QPP regression led to an overall decrease in local connectivity in the DMN and TPN and a decrease in anti-correlation between the DMN and TPN. However, regression of the Control QPP from Control scans led a greater number of functional connectivity differences (*n* = 494; 76% of all connections within and across the DMN and TPN) than regression of the ADHD QPP from ADHD scans (*n* = 280; 43% of all connections within and across the DMN and TPN). Though the overall direction of functional connectivity differences was the same, removal of the ADHD QPP from ADHD scans resulted in far fewer significant changes in functional connectivity compared to removal of the Control QPP from Control scans. A comparison of functional connectivity differences that includes the regression of the ADHD QPP from Control scans and regression of the Control QPP from ADHD scans is shown in Supplementary Figure 2.

## 4 Discussion

We studied the dynamics of BOLD fluctuations in individuals with ADHD through the investigation of quasi-periodic patterns in the brain. QPPs have been shown to contribute to functional connectivity in key functional networks and their activity has been postulated to be relevant for healthy brain function (Abbas et al., 2018a). However, until now, there has not been an investigation of QPPs in individuals with a brain disorder. ADHD is associated with atypical functional connectivity in the DMN and TPN. Given the strong involvement of the two networks in the spatiotemporal pattern captured within QPPs, we hypothesized a relationship between QPPs and the atypical functional connectivity associated with ADHD. We find that QPPs contribute to functional connectivity in the very connections that are disrupted during ADHD. Individuals with ADHD showed differences in the spatiotemporal pattern captured within the QPP, which resulted in the QPP contributing less to functional connectivity in the DMN and TPN. Our observations provide insight into the possible mechanisms behind functional connectivity differences seen in individuals with ADHD, allowing a better understanding of the etiology of the disorder.

### 4.1 Default mode and task positive networks

The brain regions in the DMN and TPN that were common to both the Control and ADHD groups largely agreed with previous literature (Raichle, 2015; Fox et al., 2005). The ROIs unique to the DMN or TPN masks acquired from the ADHD group were difficult to interpret as only a relatively small number of their voxels overlapped with the DMN or TPN masks. The differences observed could be within the margins of error associated with functional MRI or the ROI boundaries in a brain atlas such as the one used in this study. One exception to this was the inclusion of the thalamus in DMN mask from individuals with ADHD. Areas in the thalamus have previously been shown to have increased connectivity with DMN regions in individuals with ADHD (Tian et al., 2006; Tian et al., 2008), which would explain their inclusion in the ADHD DMN.

There was a consistent difference between the two groups in the strength of correlation of all DMN and TPN ROIs with the PCC, the seed used to create the masks. DMN ROIs in the ADHD group had a weaker correlation with the PCC and TPN ROIs in the ADHD group had a weaker anti-correlation with the PCC. This is likely a reflection of the observation that overall DMN/ TPN anti-correlation was also weaker in individuals with ADHD. Strong anti-correlation between the DMN and TPN is a sign of healthy brain function (Fox et al., 2005) and is related to performance on vigilance tasks (Thompson et al., 2013). Indeed, it has been previously shown that individuals with ADHD show decreased anti-correlation between DMN and TPN activity (Sripada et al., 2014). This disruption has been shown to affect task performance and pharmaceutical solutions have been suggested to alleviate the atypical functional connectivity between the two networks (Querne et al., 2014; Rubia et al., 2009b). Our observations confirm a decreased anti-correlation in individuals with ADHD. Reproducing previous findings was a critical first step in analyzing the dynamics of the BOLD signal and investigating how QPPs may be contributing to differences observed through traditional static analyses of fMRI.

### 4.2 Quasi-periodic patterns

Quasi-periodic patterns were observed in both the Control and ADHD groups. In each case, the spatiotemporal pattern captured in the QPP showed the DMN-to-TPN transition reported in previous studies (Majeed et al., 2011; Yousefi et al., 2018; Abbas et al., 2018a). The pattern lasted approximately 20 seconds and occurred quasi-periodically in the functional scans from both groups.

This was the first investigation of QPPs in individuals with a brain disorder. Functional connectivity disruptions in the DMN and TPN have been widely reported in individuals with ADHD (for reviews, see Konrad & EIckhoff, 2010; Cortese et al., 2012; Hart et al., 2012). Given the involvement of the two networks in QPPs, it was pertinent to compare the spatiotemporal pattern of the QPPs between the Control and ADHD groups. A similar spatial comparison was carried out in Abbas et al. (2018). QPPs were differentiated between resting-state and task-performing individuals. The observed differences were reflective of the working-memory task being performed. It assisted in explaining the functional connectivity changes that occur in task-performing individuals. Here, we hypothesize that any differences in the Control and ADHD QPPs may help explain the functional connectivity differences seen between the two groups.

Figure 2c and Supplementary Table 3 demonstrate that the spatiotemporal pattern was largely similar in the Control and ADHD QPPs. However, the few observed differences were telling. Table 2 lists DMN or TPN ROIs that had anti-correlated timecourses between the Control and ADHD QPPs. Among the DMN ROIs, the cingulate gyrus and precuneus were also the regions that showed decreased functional connectivity within the DMN when functional connectivity was compared in Figure 4b. Among the TPN ROIs, the the middle and inferior frontal gyri showed decreased anti-correlation with DMN regions when functional connectivity was compared. Though the differences in the spatiotemporal pattern between the Control and ADHD QPPs were small, they aligned well with the region-to-region functional connectivity differences observed between the two groups. Hence, spatiotemporal differences in the QPPs between the two groups were able to predict functional connectivity differences in individuals with ADHD.

A key difference between the two QPPs was the magnitude of the difference between the DMN and TPN timecourses, as is demonstrated in Figure 2b on the right. The anti-correlation between the DMN and TPN was stronger in the Control QPP compared to the ADHD QPP. The pattern-finding algorithm used to acquire the QPPs averages occurrence of the spatiotemporal pattern over the course of the functional timeseries. Hence, the difference in magnitude of DMN/TPN anti-correlation between the two groups’ QPPs is a reflection of a general trend in the data, rather than a consequence of the randomly-selected starting segment used to initiate the pattern-finding algorithm. This is a key difference in the QPPs acquired from the Control and ADHD groups, as it can have a strong effect on the overall contribution of QPPs to functional connectivity in the brain, discussed in the next section.

Comparison of the strength and frequency of the Control and ADHD QPPs in their respective functional scans showed no differences between the two groups (Figure 3). This is also an important observation as the different effects the two QPPs had on functional connectivity can be attributed only to the spatiotemporal differences outlined above, rather than any difference in the level of presence of QPPs in the functional scans. Figure 3 also demonstrated that the regression method used in this study was effective in significantly attenuating the presence of the QPPs in the scans. The efficacy of the regression was critical as it allowed the investigation of the contribution of QPPs to functional connectivity by essentially removing them from the BOLD signal.

### 4.3 Functional connectivity

The region-to-region functional connectivity comparisons shown in Figure 4 required strict multiple comparisons correction due to the number of hypotheses being tested. However, comparison of the distribution of functional connectivity strength within the DMN, within the TPN, and between the DMN and TPN only required testing one hypothesis each. This allowed us to conclude with confidence that the local functional connectivity within the DMN and TPN was significantly lower in individuals with ADHD. Additionally, functional connectivity analysis further demonstrated that the strength of anti-correlation between the DMN and TPN was weaker in the ADHD group. Observations of the overall differences in DMN and TPN functional connectivity between the two groups continue to align with previous reports (Konrad & Eickhoff, 2010; Weyandt et al., 2010; Sripada et al., 2014; Uddin et al., 2008).

Figures 4c and 4d show that regression of the QPP from functional scans resulted in functional connectivity differences following a similar trend in both groups. Local connectivity in the DMN and TPN was reduced and anti-correlation between the DMN and TPN was weakened. This demonstrates that QPPs play a role in maintaining the functional connectivity within and across the DMN and TPN. Earlier, we saw that functional connectivity differences in individuals with ADHD follow the same trend. Our observations suggest that QPPs help maintain typical functional connectivity in the same regions that tend to develop atypical connectivity in ADHD. Hence, it may be that the functional connectivity differences in individuals with ADHD are the result of a failure of QPPs to maintain healthy functional connectivity as they do in healthy individuals.

Figures 4c and 4d also show that the Control QPP contributes to functional connectivity within and across the DMN and TPN with far greater effect than the ADHD QPP. The number of connections that were significantly affected by regression of the Control QPP was 76% greater than the number of connections significantly affected by regression of the ADHD QPP. We know that the strength and frequency of both QPPs was similar in their respective functional scans. Hence this difference is likely a result of the spatiotemporal differences in the Control and ADHD QPPs. The difference in magnitude of anti-correlation between the DMN and TPN within the spatiotemporal pattern of the QPP (Figure 2b, *right*) suggests that QPPs are contributing to an overall smaller percentage of the spontaneous BOLD signal fluctuations in individuals with ADHD. This would reduce their contribution to functional connectivity.

Figure 4b shows the difference in functional connectivity between the Control and ADHD groups. Notably, it distinguishes between the functional connectivity differences observed between the two groups before and after regression of native QPPs from the functional scans. Both the nature and the number of functional connectivity differences are different between these two comparisons. When the original functional scans were compared, the functional connectivity differences showed partial decrease in local connectivity in the DMN and reduced anticorrelation between DMN and TPN regions. These differences follow the trend of previous reports on functional connectivity disruptions in individuals with ADHD. However, when the QPP-regressed functional scans were compared, the trend of the differences was reversed and the number of functional connectivity differences increased: When comparing QPP-regressed Control scans to QPP-regressed ADHD scans, local connectivity in the DMN and TPN and anticorrelation between regions in the DMN and TPN increased. This is most likely due to the varying effects of the Control and ADHD QPPs on functional connectivity in the DMN and TPN. When the Control QPP was regressed from Control scans, it led to a large number of functional connectivity differences, as is visible in Figure 4c. When the ADHD QPP was regressed from ADHD scans, it led to a relatively smaller number of functional connectivity differences. Hence, the difference between the two comparisons in Figure 4b is a result of the ADHD QPP failing to contribute as strongly to functional connectivity in the DMN and TPN. Interestingly, comparison of functional connectivity between the Control and ADHD groups after native QPP regression demonstrates how QPPs are contributing to functional connectivity differently in individuals with ADHD. In fact, the greater the increase in functional connectivity differences observed between the two groups after QPP regression, the more the QPP in individuals with ADHD is failing to contribute to functional connectivity. This further demonstrates the relevance of QPPs in understanding the mechanisms behind functional connectivity differences in individuals with ADHD.

### 4.4 Implications for ADHD

The static functional connectivity differences in the ADHD group observed in our analysis have largely been reported in previous literature. Functional connectivity in the DMN has been shown to be decreased in individuals with ADHD (Rubia et al., 2007; Uddin et al., 2008; Liddle et al., 2010; Wilson et al., 2011; Yu-Feng et al., 2006). Reviews of several fMRI studies on ADHD (Cortese et al., 2012; Hart et al., 2012) have revealed a consistent decrease in BOLD activation in attentional networks, loosely similar to the TPN investigated in this study. Studies have also shown decreased activation in attentional networks similar to the TPN during task-based fMRI scans (Schneider et al., 2010; Rubia et al., 2009b). Increase in functional connectivity between brain regions in the DMN and TPN, which we refer to instead as a decreases in DMN/TPN anticorrelation, has also been reported (Hoekzema et al., 2013; Konrad & Eickhoff, 2010)

Analysis of the dynamics of the BOLD signal have allowed researchers to understand the mechanisms behind functional connectivity differences seen in individuals with other brain disorders (Sakoglu et al., 2010; Damaraju et al., 2012; Damaraju et al., 2014; Jones et al., 2012; Holtzheimer & Mayberg, 2011). For example, Sakoğlu et al. (2010) and Damaraju et al. (2012; 2014) demonstrated real-time inter-network interactions being disrupted during an auditory oddball task in individuals with Schizophrenia and the relative rigidity of time-varying network functional connectivity compared to healthy controls. Jones et al. (2012) showed that static functional connectivity differences observable in individuals with Alzheimer’s Disease may exist due to certain dominant sub-network configurations of the brain’s default mode network, only observable through dynamic analysis. Similarly, Holtzheimer and Mayberg (2011) argue that functional connectivity differences seen in individuals with Major Depressive Disorder are due to a tendency of network activity to linger in ‘down states’ longer compared relative to healthy controls, indiscernible through a static analysis of functional connectivity. However, analyses of the dynamics of BOLD signal in individuals with ADHD has been limited (Durston et al., 2003).

Studies sensitive to the time-varying changes in BOLD in individuals with ADHD have mostly focused on task-based BOLD activation in relevant brain regions (Schneider et al., 2010; Rubia et al., 2009b; Liddle et al., 2011; Yang et al., 2011; Siqueira et al., 2014). Sonuga-Barke and Castellanos (2007) showed that in the context of pathological conditions, the dynamics of the default mode network can affect attentional control in individuals. Outside the context of ADHD, Thompson et al. (2013) demonstrated that the dynamics of DMN and TPN activity can predict vigilance in performance on a psychomotor vigilance task. Given that QPPs can be used to study the dynamics of DMN activity in both resting-state and task-performing individuals, they have the potential to provide insight into the static functional connectivity differences observed in individuals with ADHD. We find that this is indeed the case. Since analyses focused on QPPs consider the time-varying component of BOLD signal, they can provide a more sensitive analysis of differences in individuals with ADHD. For example, the number of region-to-region functional connectivity differences observed between the Control and ADHD groups was small. However, when the same comparison was done after regression of the QPP, the number of functional connectivity differences between the two groups was appreciably larger.

It has been demonstrated that using static functional connectivity differences as a biomarker in individuals with ADHD is not yet the most accurate way to differentiate them from healthy controls: Brown et al. (2012) showed that personal characteristic data—such as age, gender, and performance on different IQ tests--was more accurate in predicting ADHD diagnosis than static functional connectivity differences. Analysis techniques that focus on the dynamics of the BOLD signal, such as the one shown in this study, may provide greater sensitivity to differences in individuals with ADHD. The functional connectivity analysis presented in Figure 4 shows a greater number of differences between groups compared to traditional methods, which may provide a more sensitive prediction of ADHD diagnosis. This introduces the possibility of using disruptions in QPPs as a potential biomarker of disease.

It is important to note that the results from this study do not address a critical question: Is the disruption in the QPPs causing the functional connectivity differences seen in ADHD, or is ADHD causing the disruption seen in the QPPs? However, it has been suggested—and preliminary experiments in rodents show--that QPPs have a neurophysiological driver in deep brain nuclei. Induced disruption of locus coeruleus input through injection of DSP-4 (N-(2-chloroethyl)-N-ethyl-2-bromobenzylamine) resulted in a dramatic attenuation of QPP activity in anesthetized rodents (Abbas et al., 2018b). If this is indeed the case, a hypothesized pathway of the etiology of ADHD would link initial disruptions in deep brain nuclei with abnormalities in the spatiotemporal pattern of QPPs, resulting in the functional connectivity differences seen in individuals with ADHD.

Local field potential (LFP) electrophysiological recordings in anesthetized rats conducted simultaneously with fMRI have shown that QPPs are correlated with infra-slow electrical activity (Pan et al., 2013). Infra-slow electrical activity is disrupted in individuals with ADHD: Helps et al. (2010) showed reduced attenuation of electroencephalography (EEG) power at infra-slow frequency bands (0.02-0.2Hz) in individuals with ADHD, which was associated with poor performance on attentional tasks. Monto et al. (2008) also showed that psychophysical performance is related to infra-slow fluctuations in electrical activity measured through EEG. Future investigations of the relationship between QPPs, and functional connectivity, and electrical activity could enhance the understanding of the etiology of ADHD.

### 4.5 Limitations

The dataset used in our analysis included scans collected at different facilities using different scanners and slightly different scan parameters. This has the potential to increase variability in the functional data, reducing the likelihood of observation of subtle differences between groups. However, the heterogeneity in the data also speaks to the robustness of the differences that *were* observed in individuals with ADHD. Second, though the Control group had an even distribution of males and females, the ADHD group was dominated (91%) by males. This is a reflection of the relatively higher clinical referral of boys when symptoms for ADHD are present, the existing bias in the ADHD literature towards male participants, and the tendency for females to be diagnosed with the Inattentive sub-type of ADHD, which was not used this study (Biederman et al., 2002; Arnold, 1996; Gaub & Carlson, 1997; Sharp et al., 1999). Third, the selection of only the Combined sub-type of ADHD may have helped reduce variability in the results and made analysis more straightforward, but it may have also resulted in certain differences in individuals with other types of ADHD being ignored. However, given the dramatic effect of QPPs on functional connectivity in most regions in the DMN and TPN, we believe that separate analysis of different sub-types of ADHD may not have resulted in conclusions dramatically different than the ones presented in this study, as the overall trend of DMN/TPN functional connectivity differences would have been the same.

The regression method used to ‘remove’ the QPPs from the functional data inherently assumes that QPPs are an additive component to the remaining BOLD signal. This is also addressed in Abbas et al., (2018), which studied the contributions of QPPs to functional connectivity throughout the brain in healthy adults using the same regression method. The assumption is based on multi-modal experiments in rodents that support the notion that QPPs are additive to the BOLD signal (Thompson et al., 2014). Though further work with neural recordings in animal models is required to provide ‘ground truth’ comparisons, treating QPPs as an additive signal is a reasonable first approximation.

There were multiple justifications for consolidating the 273 ROIs from the Brainnetome atlas into the 36 ROIs that were used to construct the functional connectivity matrices. Most importantly, QPPs have been shown to mainly contribute to functional connectivity in the DMN and TPN (Abbas et al., 2018a), which is also where most functional connectivity disruptions relevant to ADHD have been reported (Konrad & Eickhoff, 2010). Hence, a focus on DMN and TPN connectivity was appropriate when studying the relationship between QPPs and ADHD. Notably, only voxels in the ROIs that overlapped with the DMN or TPN masks were used, allowing the functional connectivity analysis to be specific to the two networks. Additionally, consolidation of ROIs into larger brain regions helped alleviate variability in the ROI timecourses, providing more reliable results. Finally, consolidation of the ROIs meant that the number of comparisons being performed to determine the statistical significance of a change in connectivity was reduced from 37264 to 648; a 98% decrease.

Finally, it is important to comment on the use of global signal regression during the preprocessing of all functional scans. It has been shown that global signal regression reduces variability in QPPs acquired from different individuals. In Yousefi et al. (2018), individuals were divided into two groups; those with low levels of global signal fluctuation and those with high levels of global signal fluctuation. Individuals with low levels of global signal fluctuation showed the same anti-correlated behavior of the DMN and TPN reported in this study. Individuals with high levels of global signal fluctuation showed that the global signal had an additive effect on the QPP: Though the observed spatiotemporal pattern and its frequency of occurrence was relatively unchanged, the whole-brain global changes in BOLD obscured the underlying pattern. When global signal regression was conducted in the individuals with high levels of global signal fluctuation, their QPPs aligned with those of individuals with low levels of global signal fluctuation. A primary aim of this study was to understand the effects of QPP regression on functional connectivity in the brain. If global signal had not been regressed from the functional scans, it could have served as a confounding factor in the subsequent analysis. Depending on the levels of global signal fluctuation in each individual, the spatiotemporal pattern observed in QPPs would have varied and their regression would have affected static functional connectivity differently across individuals. Hence, for a study investigating the effect of QPP regression on functional connectivity, global signal regression in all functional scans was appropriate, especially given that there are several studies already demonstrating the effects of global signal regression on functional connectivity (Murphy & Fox, 2007).

## 5 Conclusions

We confirm that functional connectivity within and across the default mode and task positive networks is disrupted in individuals with ADHD. Investigation of quasi-periodic patterns is an effective way to understand the dynamics of the BOLD signal underlying those functional connectivity differences. We find that QPPs help maintain connectivity in the same brain regions affected during ADHD. Disruptions in the spatiotemporal pattern of the QPPs may be leading to an inability of the QPPs to maintain healthy functional connectivity in those regions. This could potentially underlie the functional connectivity differences seen in individuals with ADHD and provide a more accurate understanding of the etiology of the neurodevelopmental disorder.

## Acknowledgements

The authors would like to thank Helen Mayberg, Eric Schumacher, Michael Borich, Paul Garcia, and David Weinshenker for their comments regarding the work presented here. Funding was provided by NSF BCS INSPIRE 1533260, NIH R01MH111416, and NIH R01NS078095. The data used in this study in its entirety was acquired from the ADHD 200 Sample within the 1000 Functional Connectome Project. The authors would like to acknowledge the individuals involved in collection of all data and the various funding sources that supported the NeuroImage Sample, New York University dataset, and Peking University dataset, which are included but not limited to NIMH (R01MH083246), Autism Speaks, The Stavros Niarchos Foundation, Leon Levy Foundation, an endowment provided by Phyllis Green and Randolph Cōwen, Commonwealth Sciences Foundation, Ministry of Health, China (200802073), National Foundation, Ministry of Science and Technology, China (2007BAI17B03), National Natural Sciences Foundation, China (30970802), Funds for International Cooperation of the National Natural Science Foundation of China (81020108022), National Natural Science Foundation of China (8100059), and the Open Research Fund of the State Key Laboratory of Cognitive Neuroscience and Learning.

## Supplementary Information

**Supplementary Figure 1.**
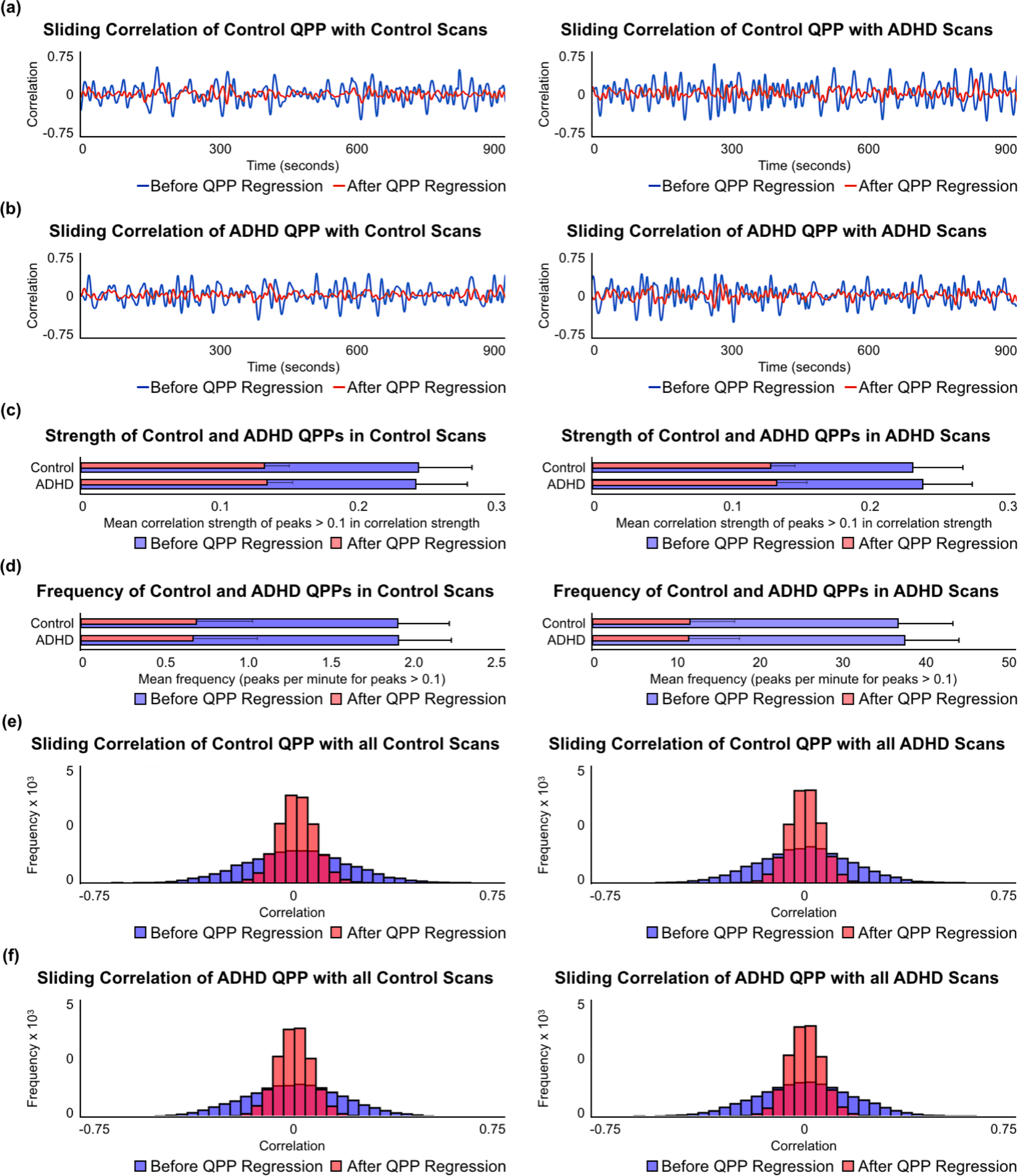
Comparison of the strength and frequency of QPPs between the Control and ADHD groups before and after removal of QPPs. This is a partial repetition of Figure 3 in the main text. However, here we compare the strength and frequency of QPPs in their non-native scans. **(a)** Example sliding correlation vectors of the Control QPP with three randomly-selected concatenated functional scans from the Control group (*left*) and the ADHD group (*right*) before (*blue*) and after (*red*) regression of the Control QPP. **(b)** Example sliding correlation vectors of the ADHD QPP with three randomly-selected concatenated functional scans from the Control group (*left*) and the ADHD group (*right*) before (*blue*) and after (*red*) regression of the Control QPP. **(c)** Mean correlation strength of peaks > 0.1 in the cumulative sliding correlation of the Control and ADHD QPPs with all Control scans (*left*) and all ADHD scans (*right*) before QPP removal (*blue*) and after QPP removal (red). **(d)** Mean frequency (peaks per minute) of peaks with correlation strength > 0.1 in the cumulative sliding correlation of the Control and ADHD QPPs with all Control scans (*left*) and all ADHD scans (*right*) before QPP removal (*blue*) and after QPP removal (red). **(e)** Histogram of the cumulative sliding correlation of the Control QPP with all Control scans (*left*) and all ADHD scans (*right*) before QPP removal (*blue*) and after QPP removal (red). **(f)** Histogram of the cumulative sliding correlation of the ADHD QPP with all Control scans (*left*) and all ADHD scans (*right*) before QPP removal (*blue*) and after QPP removal (red). There were no significant differences in strength and frequency of either QPPs in each of the groups.s

**Supplementary Figure 2.**
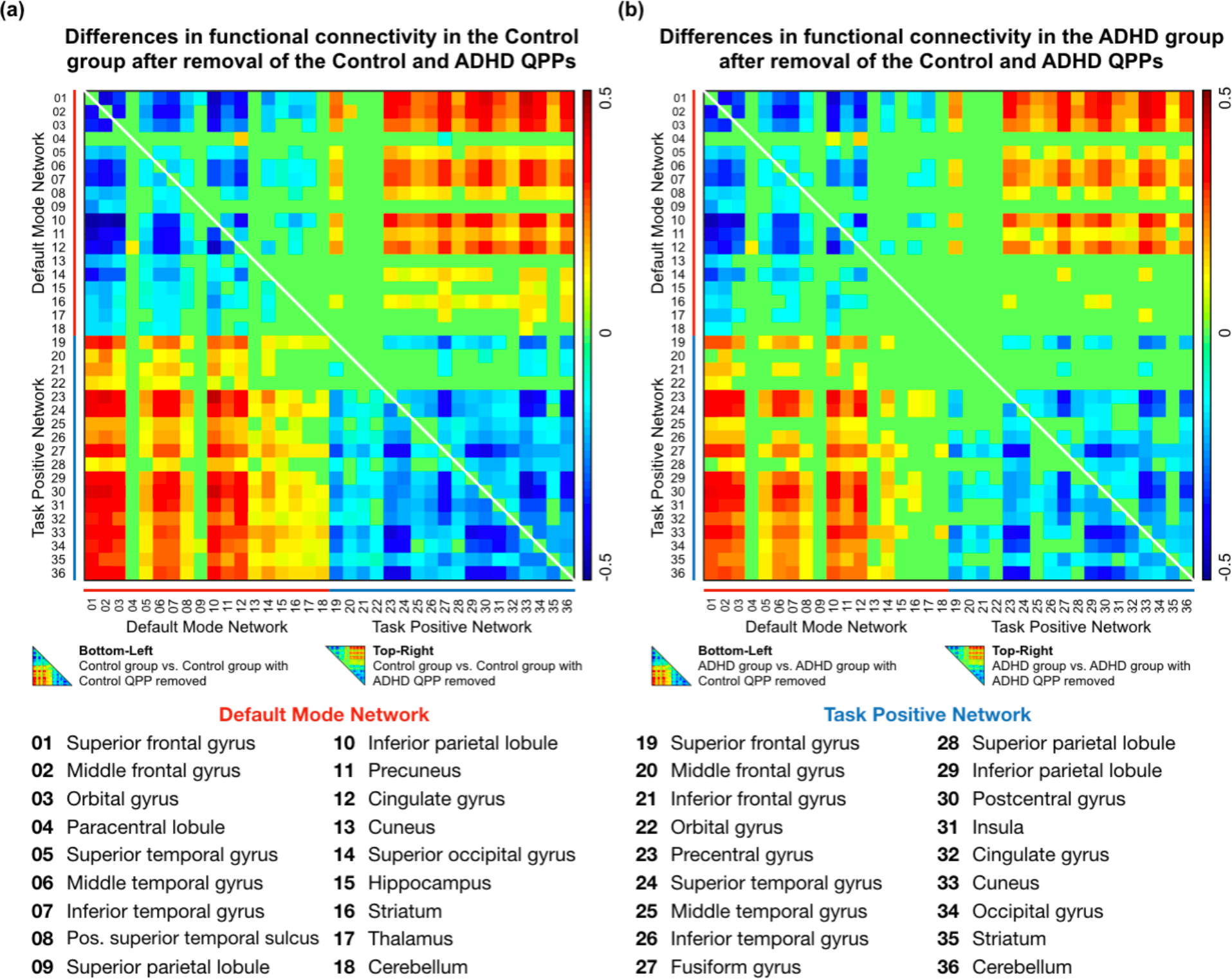
Functional connectivity in 36 ROIs within the DMN and TPN after regression of QPPs. Parts of these matrices have also been shown in Figure 4 in the main text. However, here we show the effects of regressing non-native QPPs from the functional scans in each group. **(a)** Differences in functional connectivity in the Control scans after regression of the Control (*bottom-left*) and ADHD (*top-right*) QPPs. **(b)** Differences in functional connectivity in the ADHD scans after regression of the Control (*bottom-left*) and ADHD (*top-right*) QPPs. When the Control QPP was removed from the Control data, there were 494 significant changes in functional connectivity within and across the DMN and TPN. When the ADHD QPP was removed from the Control data, there were 362 significant changes in functional connectivity. When the Control QPP was removed from the ADHD group, there were 361 significant changes in functional connectivity. Lastly, when the ADHD QPP was removed from the ADHD scans, there were only 280 significant changes in functional connectivity.

**Supplementary Table 1.**
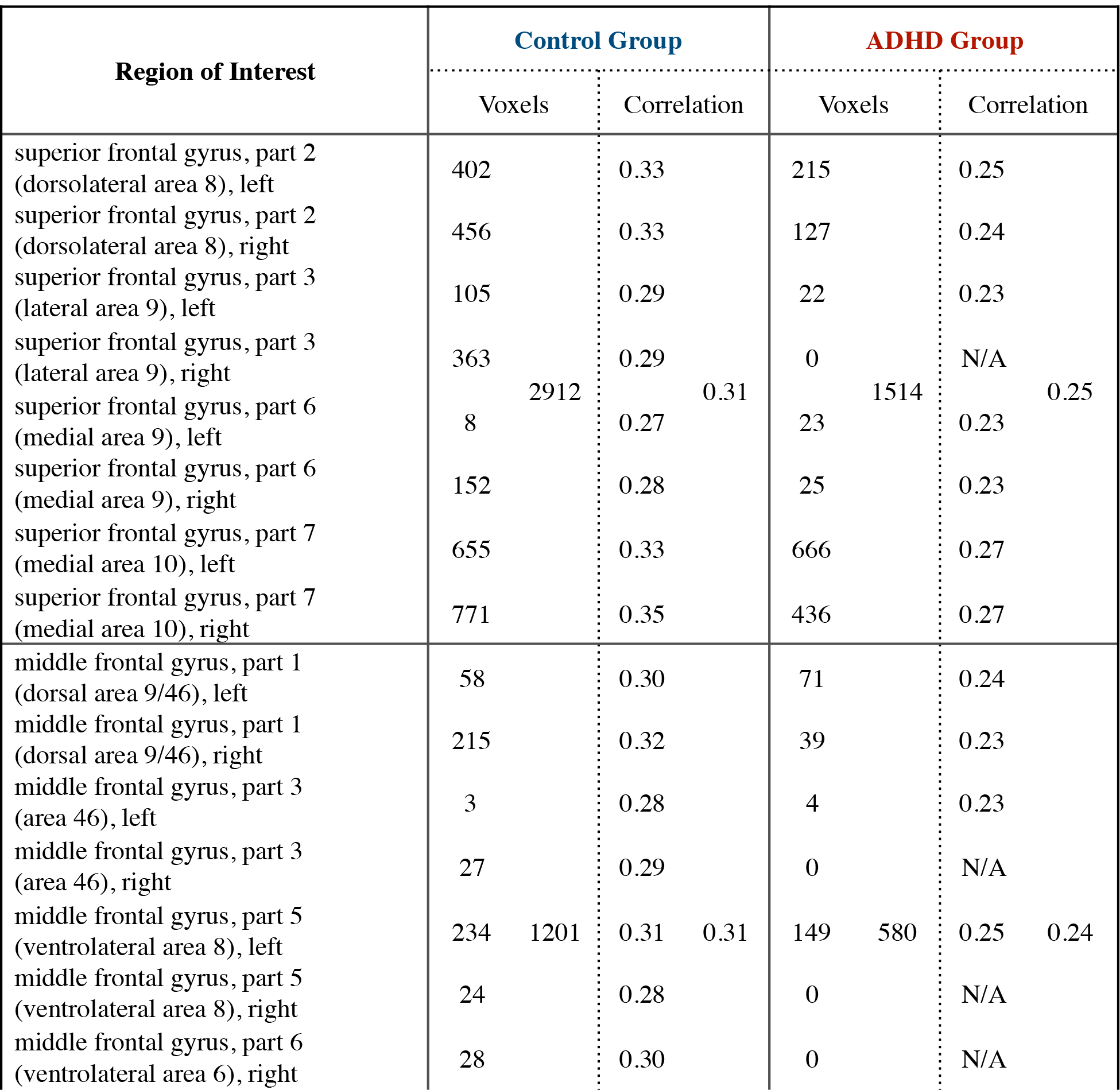
Regions of interest from the 273 ROI Brainnetome atlas from Fan et al. (2016) that were in the default mode network in control individuals and individuals with ADHD. The table shows the number of voxels in each ROI that overlapped with the DMN mask as well as how many voxels ended up in the brain regions that made up the consolidated DMN atlas. Additionally, the table shows the mean correlation with the PCC with each ROI as well as the brain regions that made up the consolidated DMN atlas.

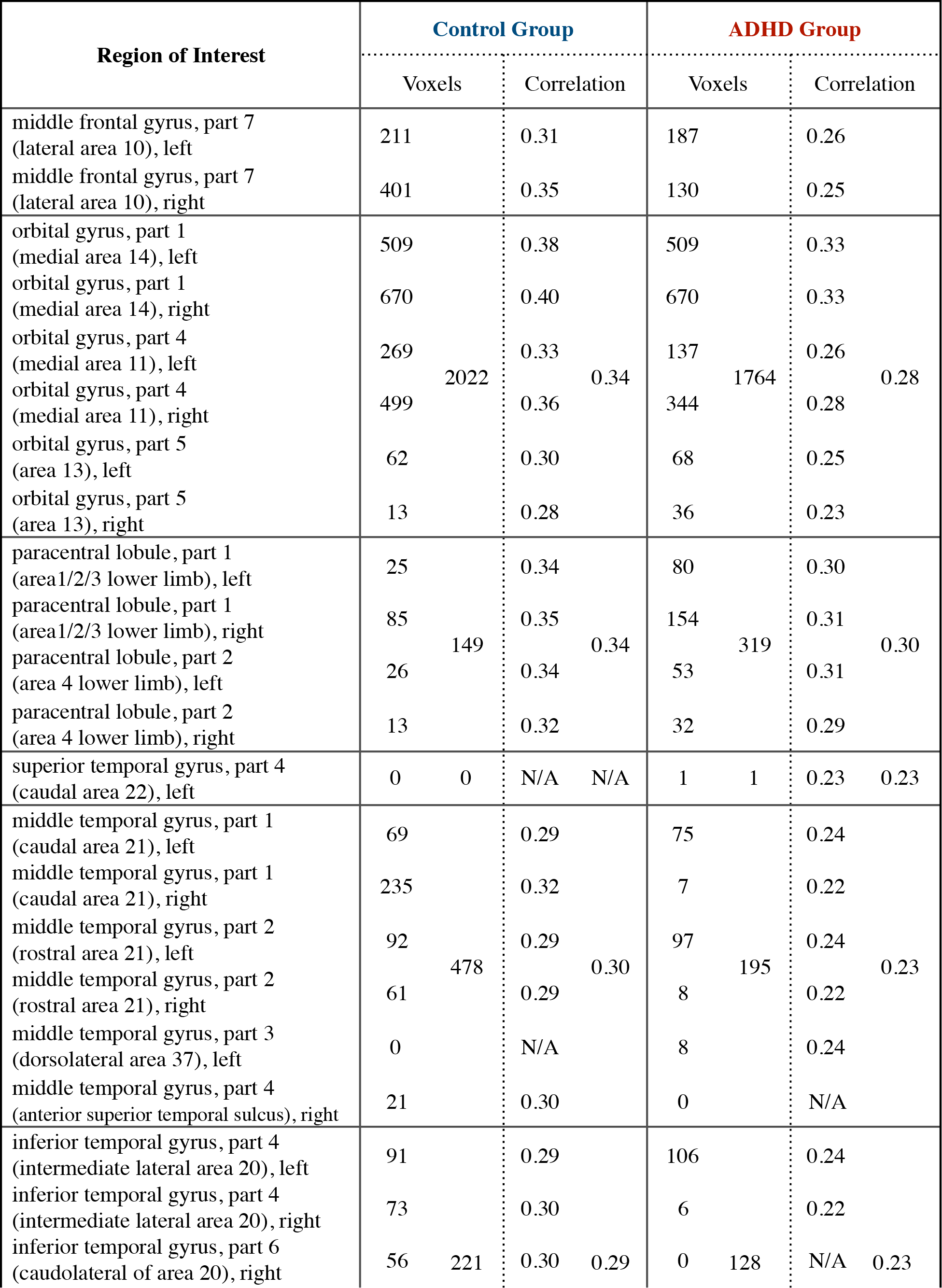

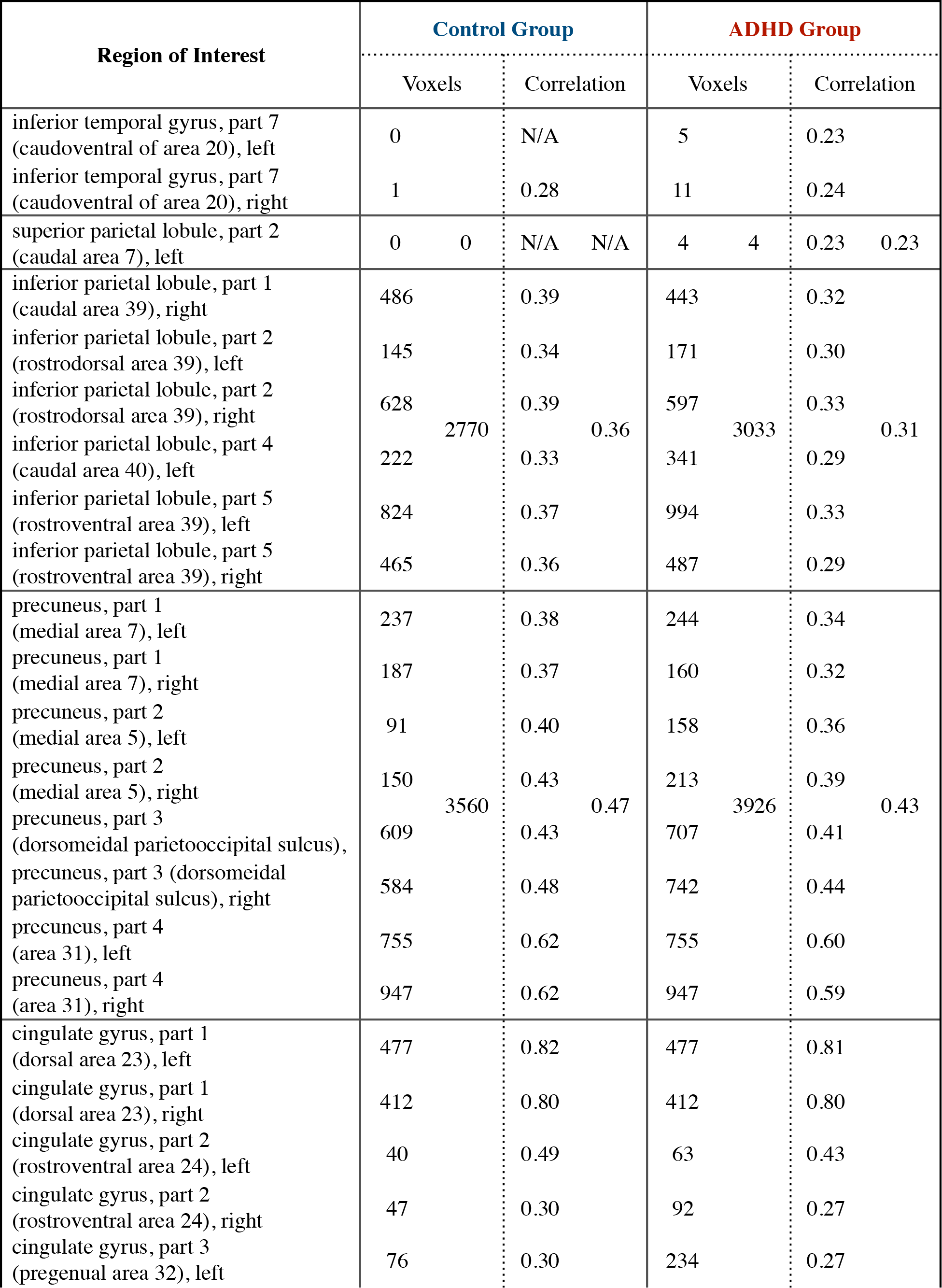

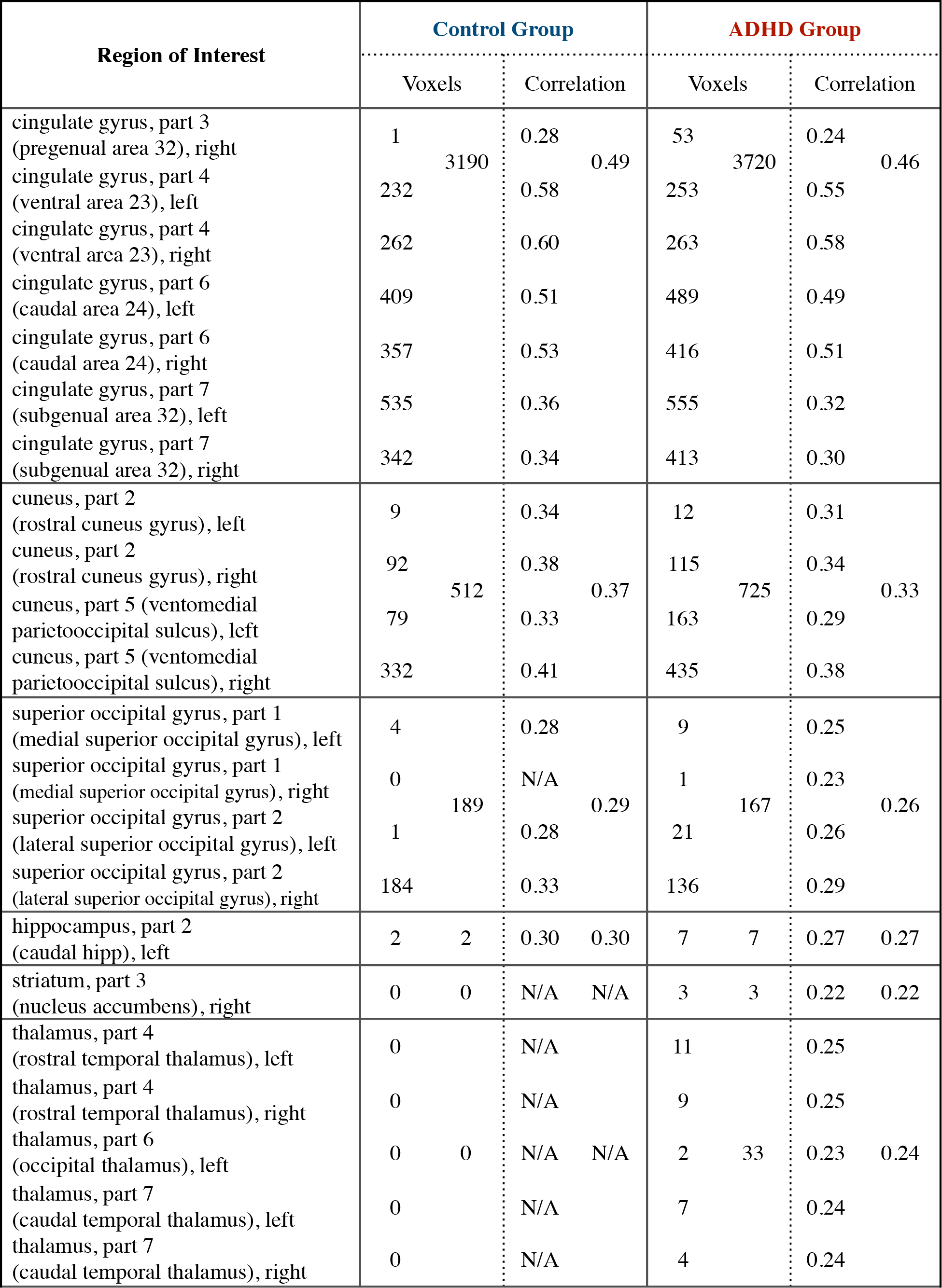

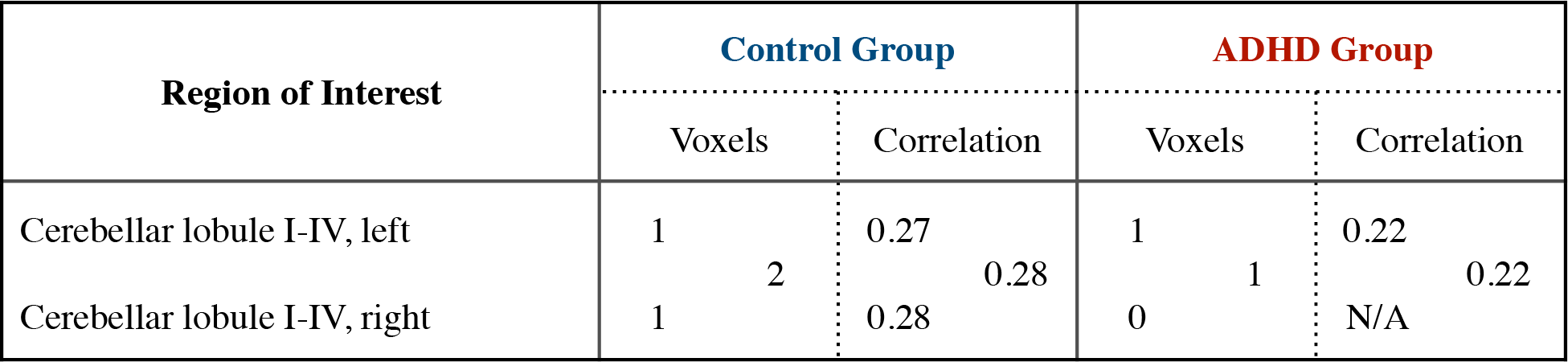

**Supplementary Table 2.**
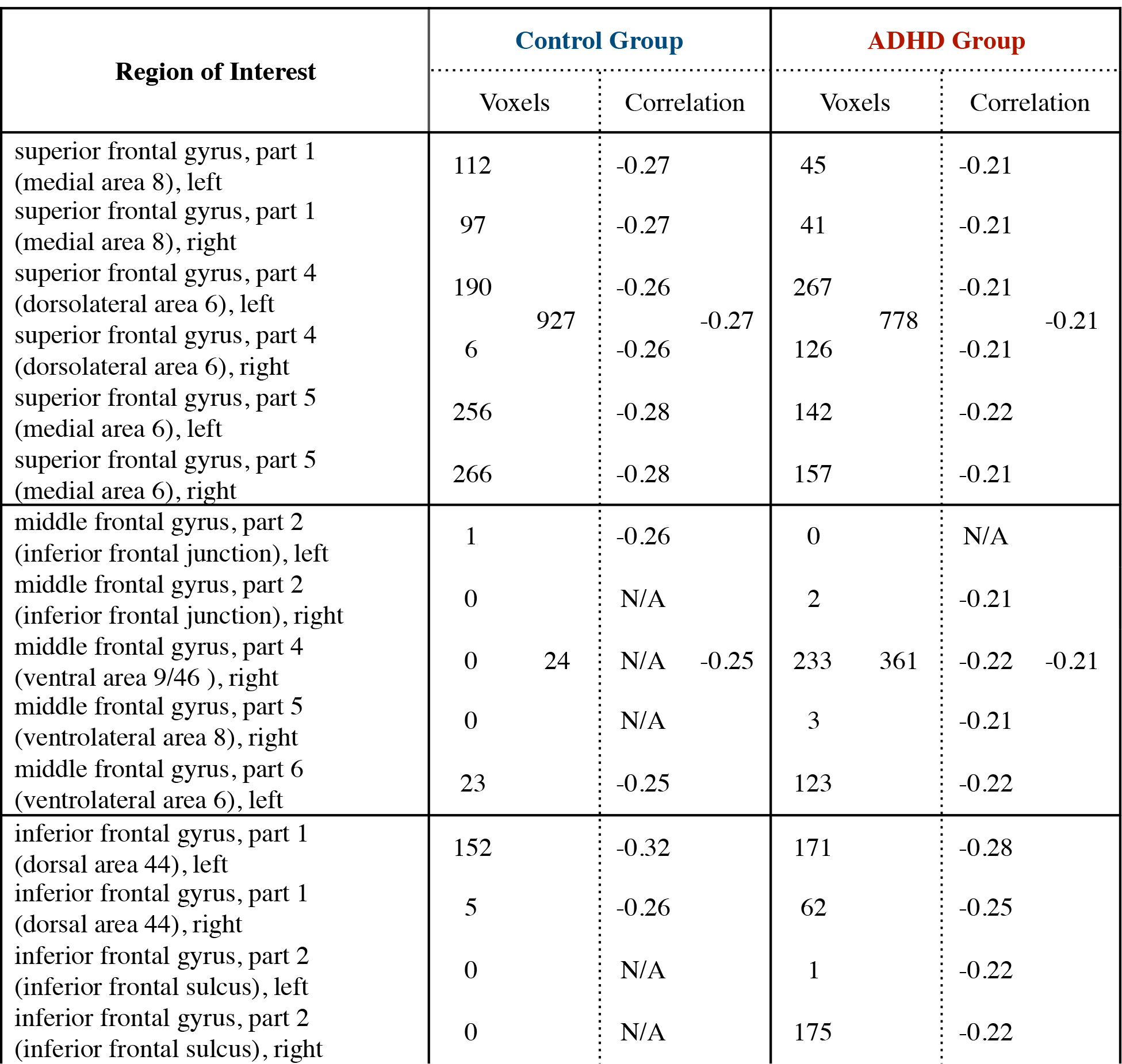
Regions of interest from the 273 ROI Brainnetome atlas from Fan et al. (2016) that were in the task positive network in control individuals and individuals with ADHD. The table shows the number of voxels in each ROI that overlapped with the TPN mask as well as how many voxels ended up in the brain regions that made up the consolidated TPN atlas. Additionally, the table shows the mean correlation with the PCC with each ROI as well as the brain regions that made up the consolidated TPN atlas.

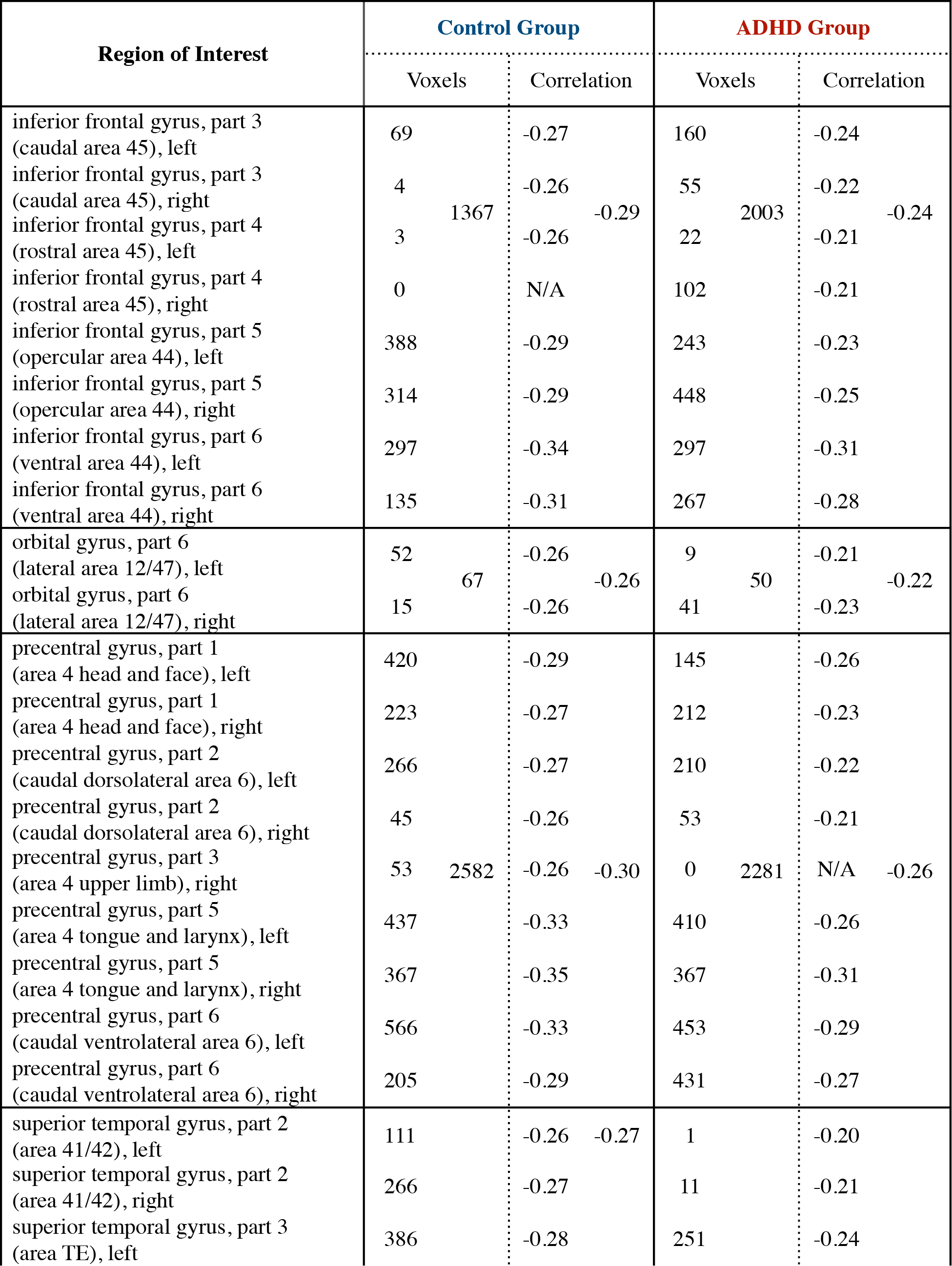

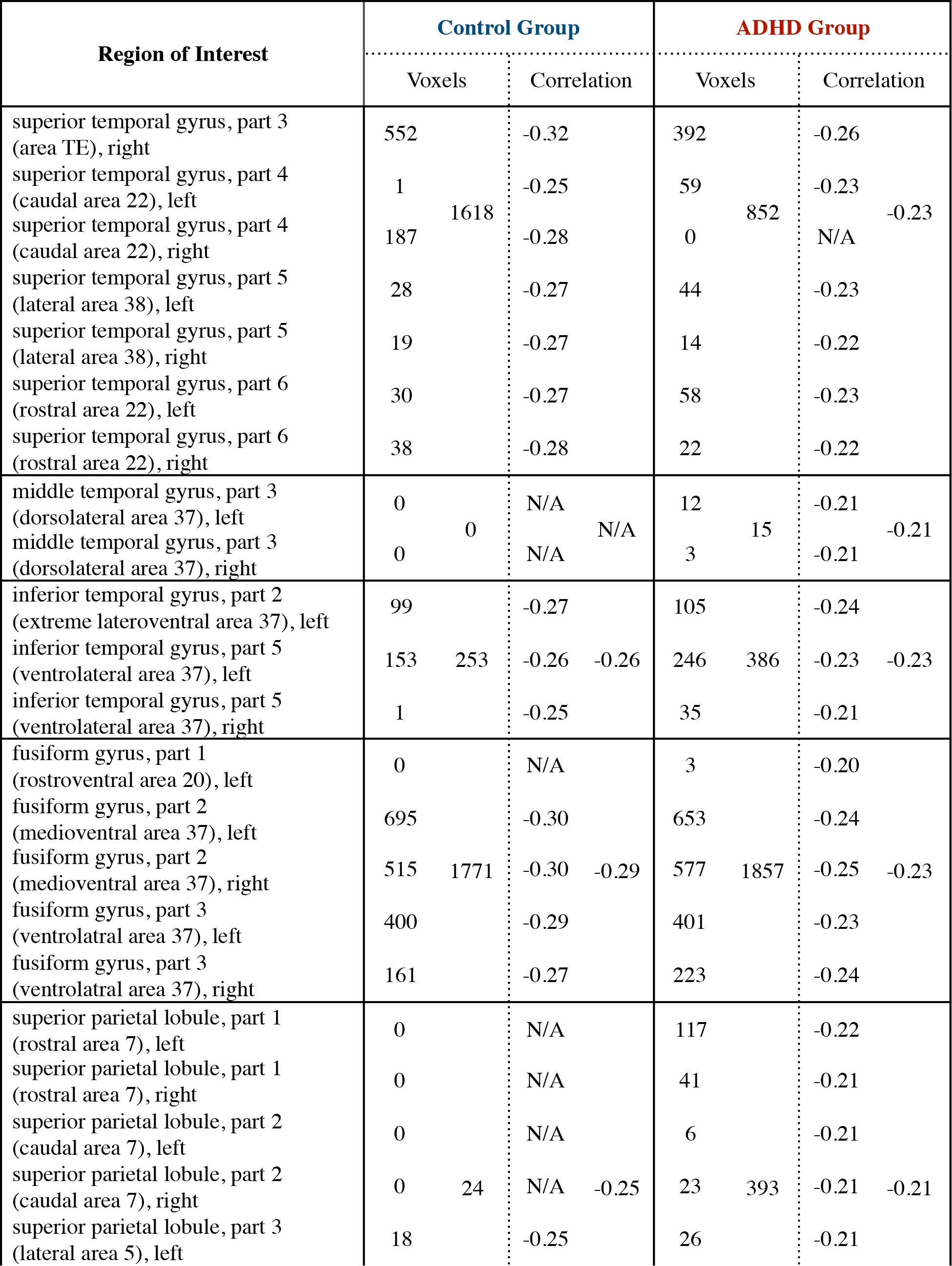

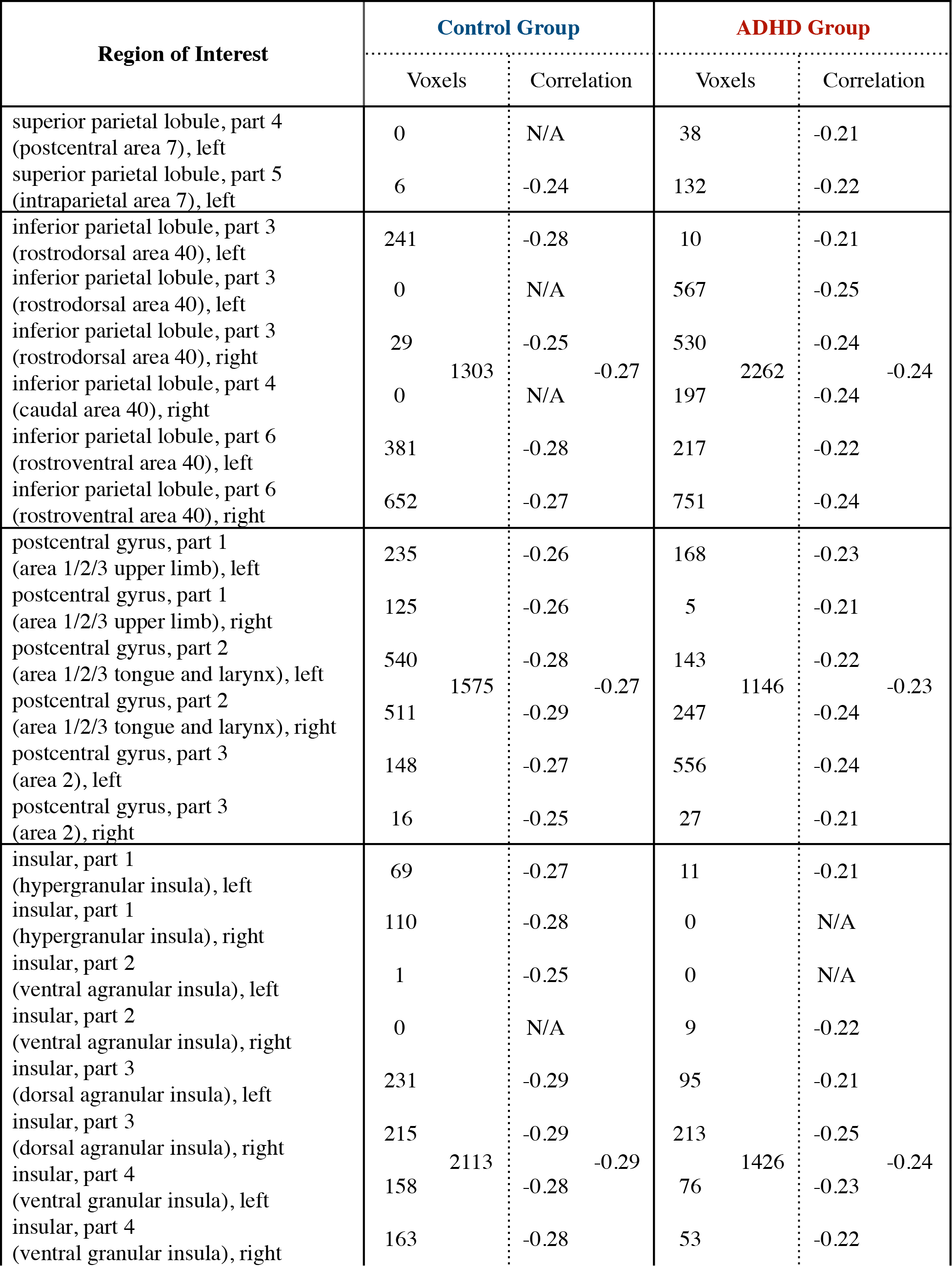

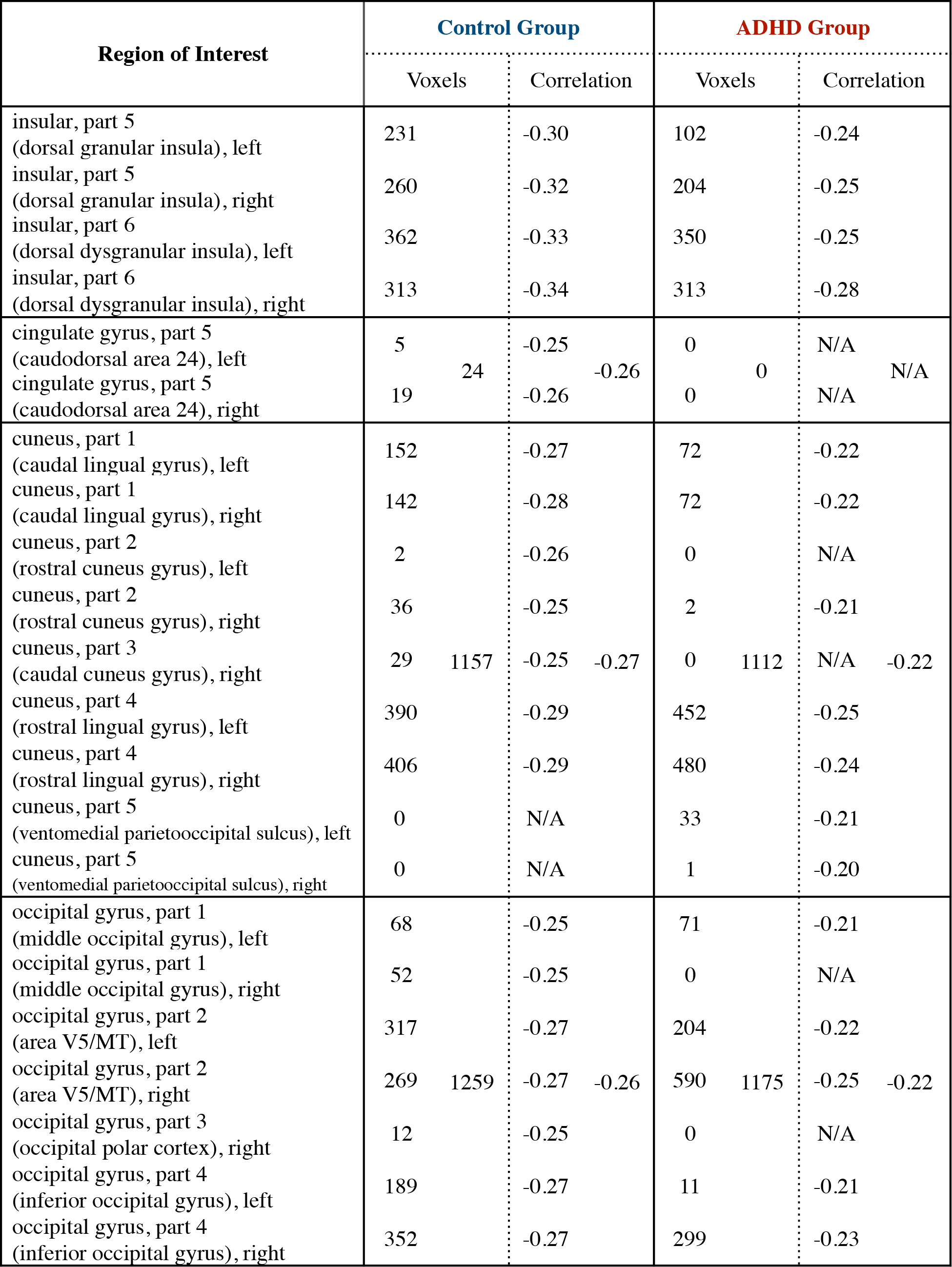

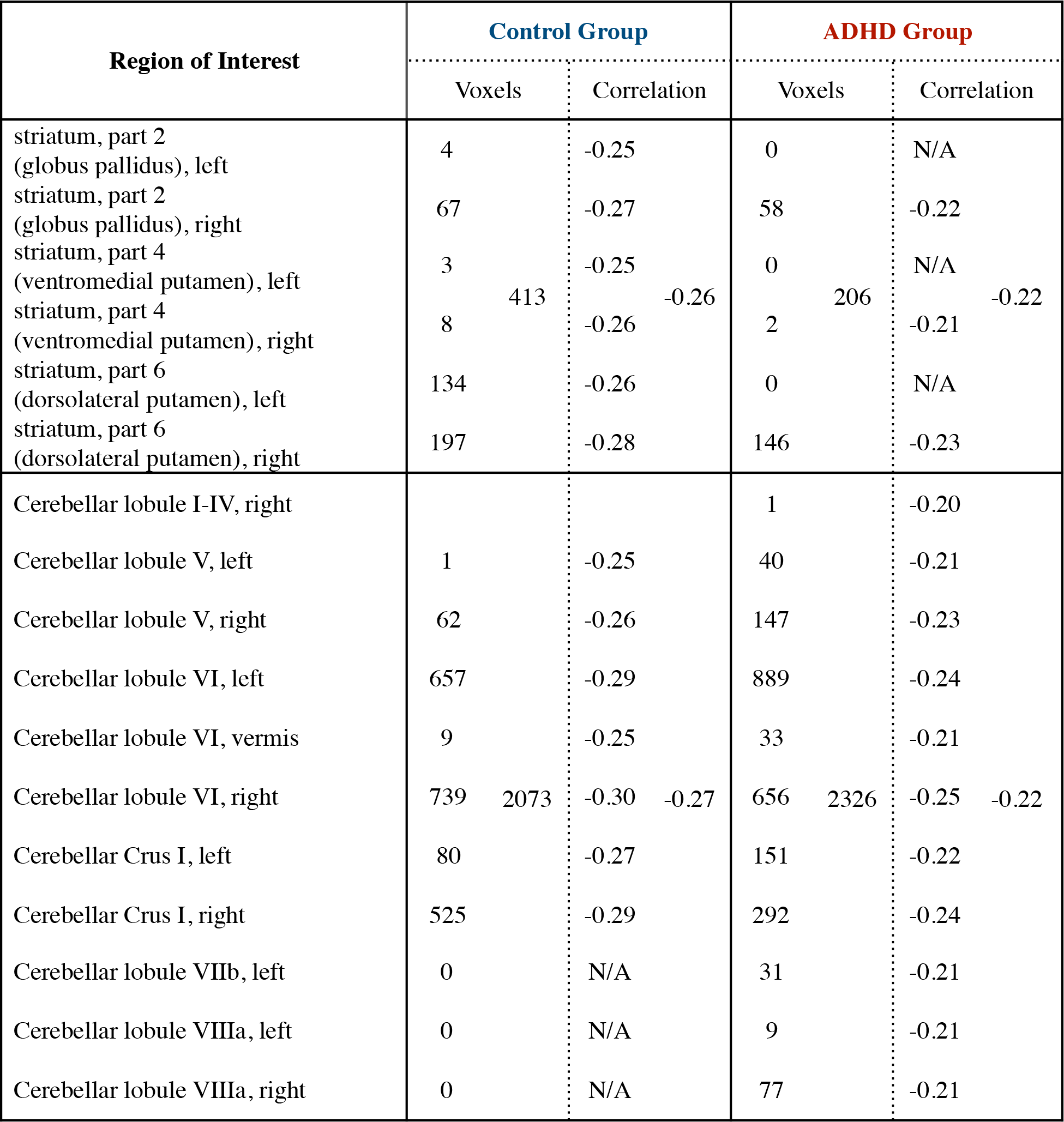

**Supplementary Table 3.**
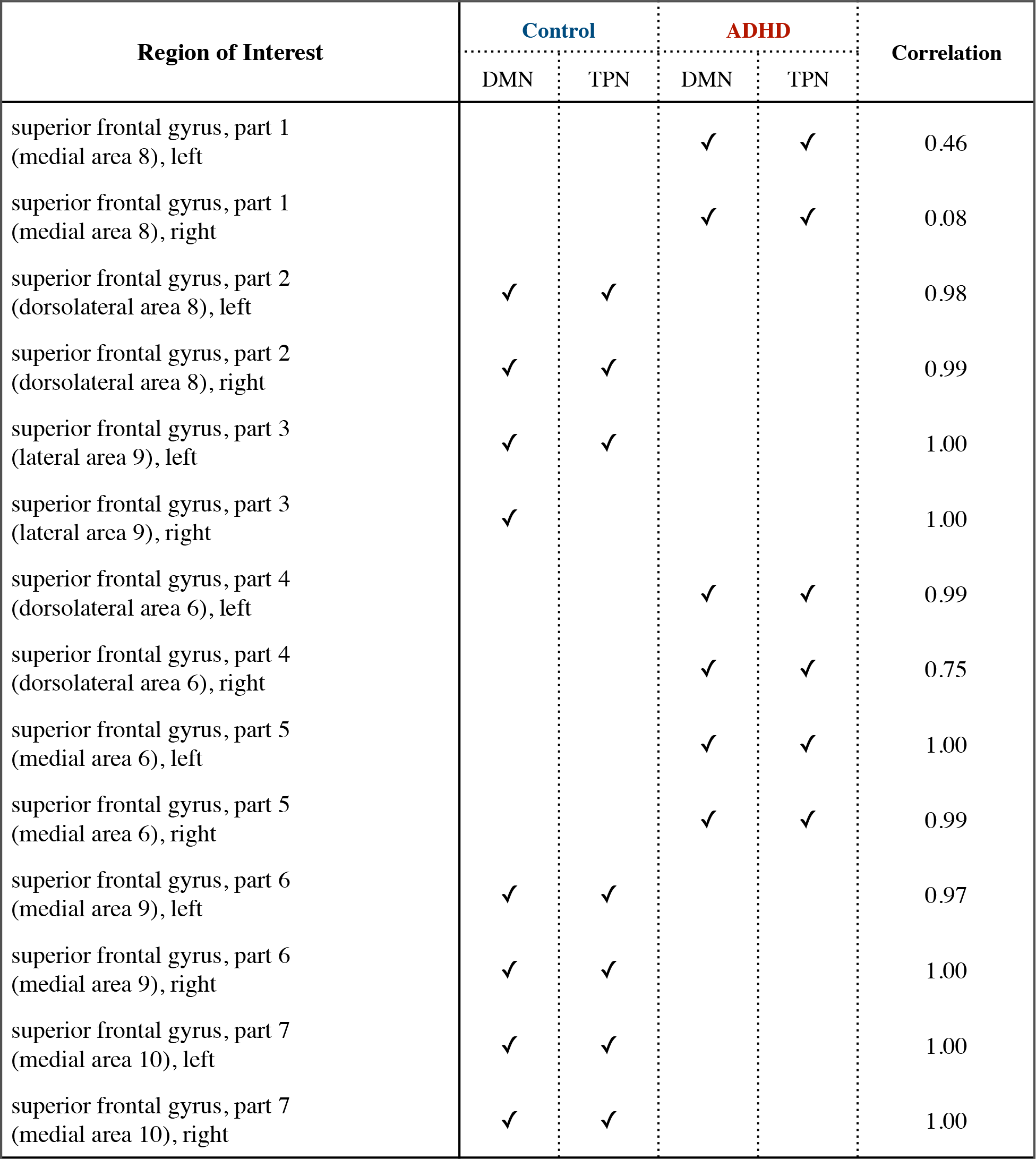
A list of regions of interest in the Brainnetome ROI atlas, whether they were included in the Control or ADHD default mode or task positive networks, and their correlation between the Control and ADHD quasi-periodic patterns. Among the four sub-columns for Correlation, the first shows the correlation value for each ROI, the second shows the mean correlation for all ROIs that were in the collective DMN from both groups, the third shows the mean correlation for all ROIs that were in the collective TPN from both groups, and the fourth shows the mean correlation for all ROIS in the respective anatomical region of the brain.

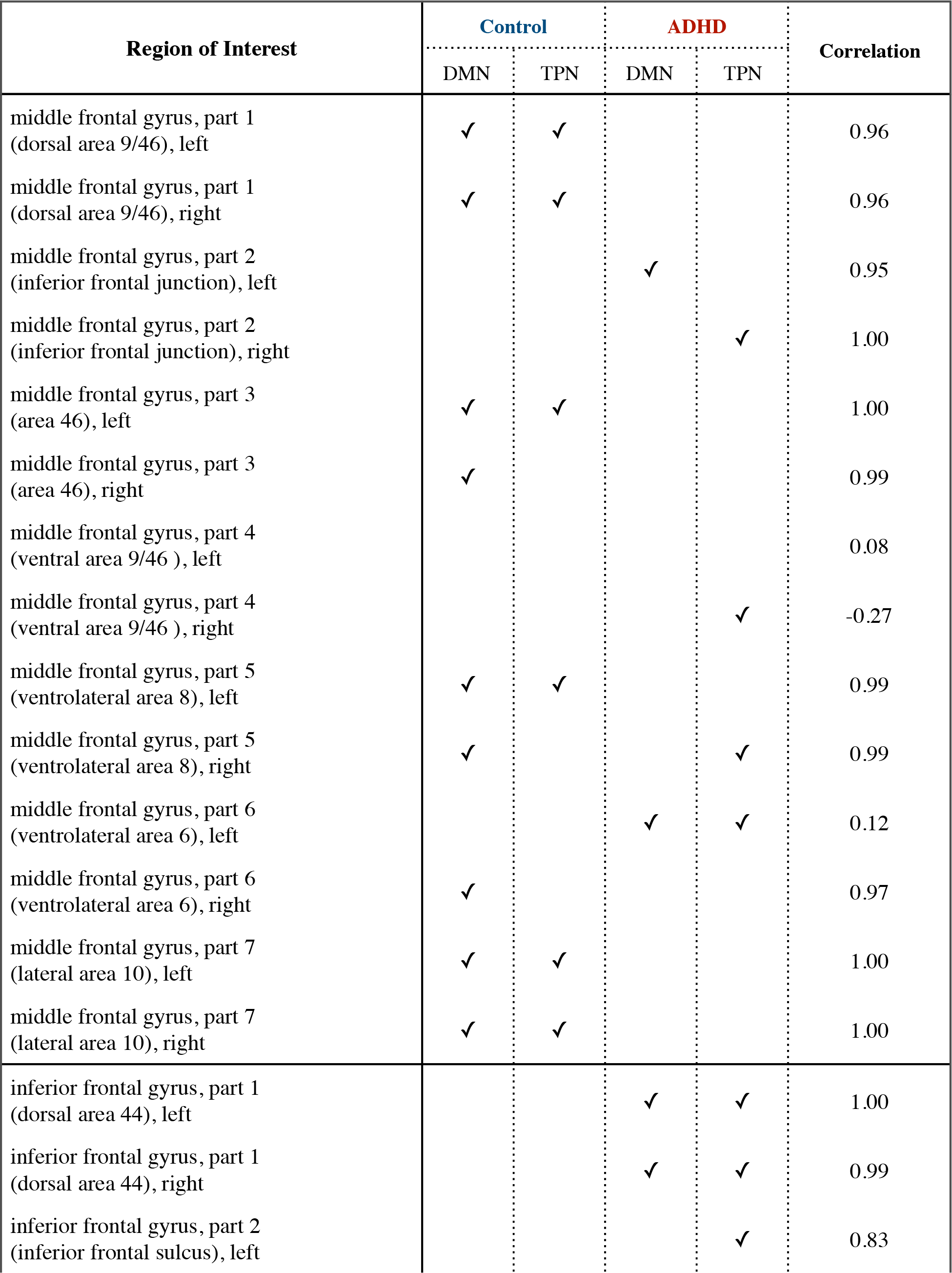

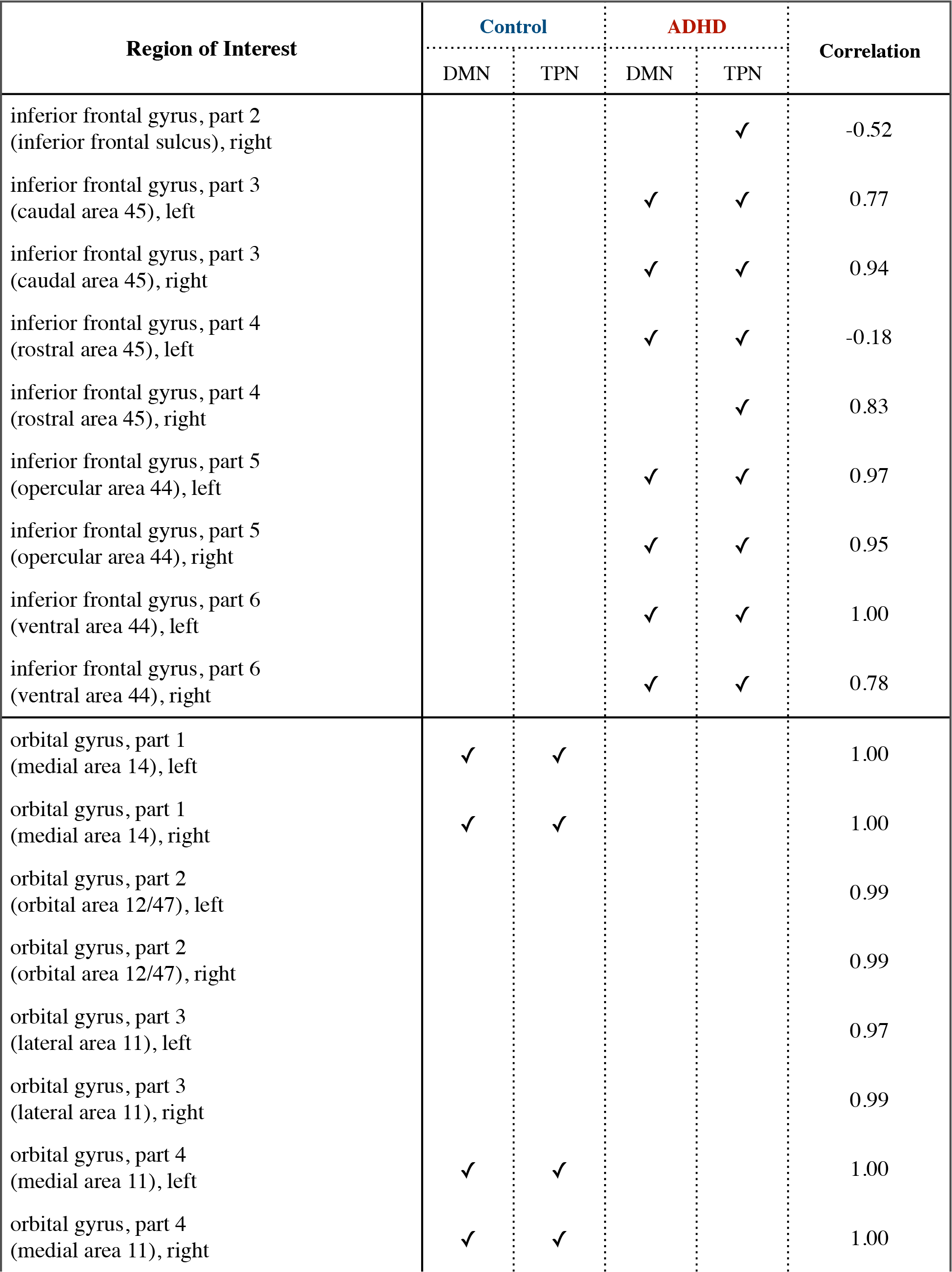

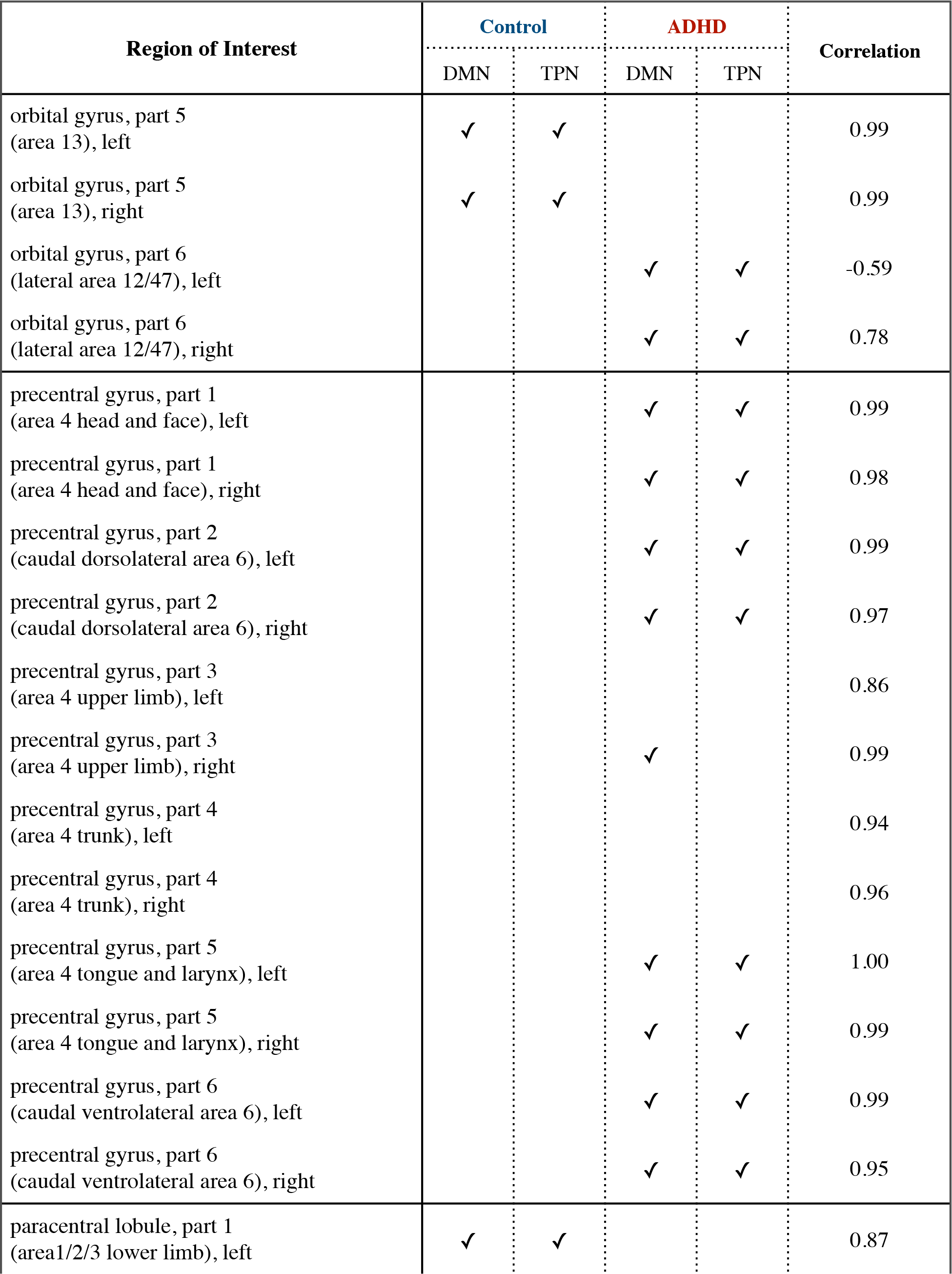

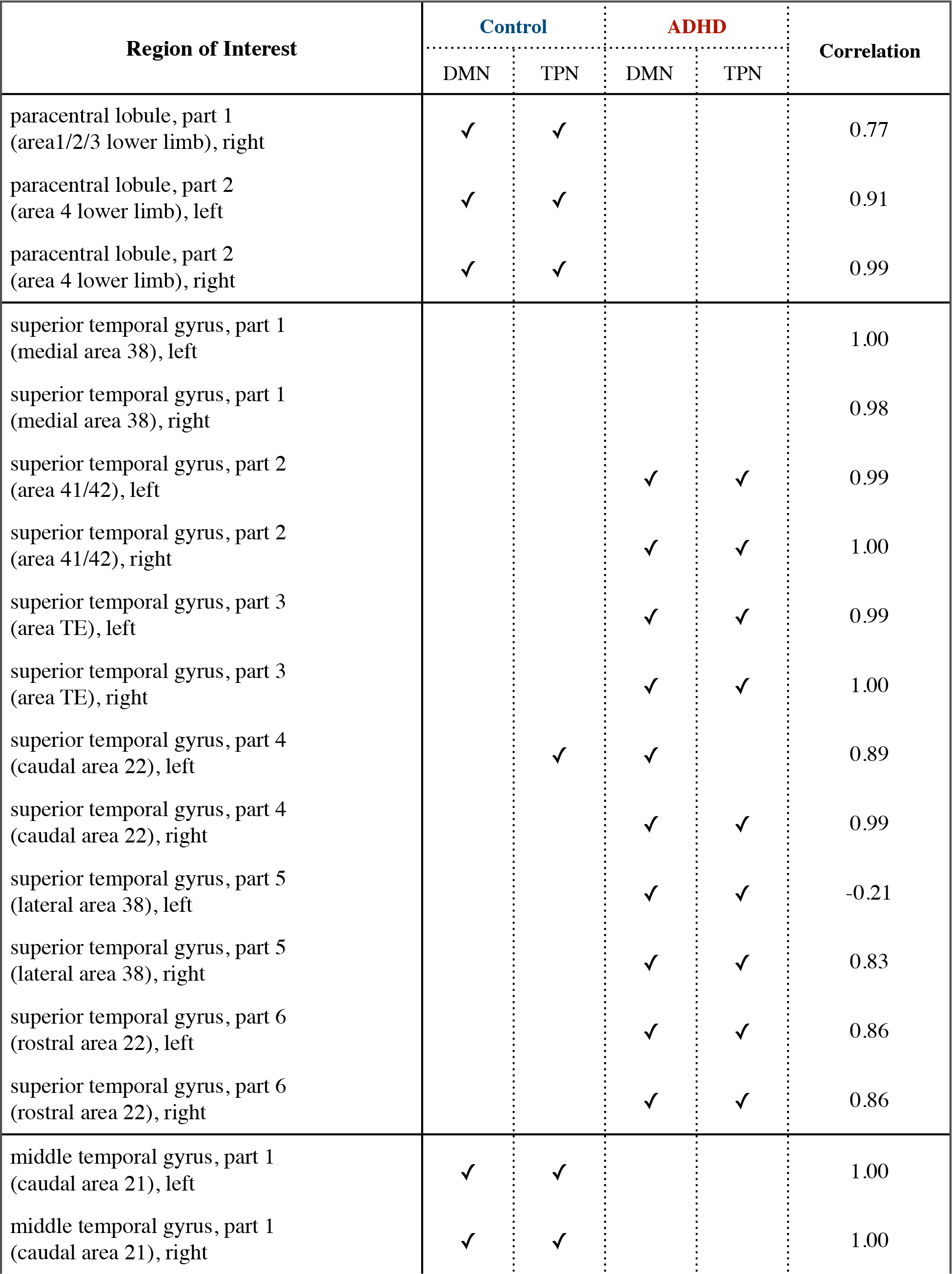

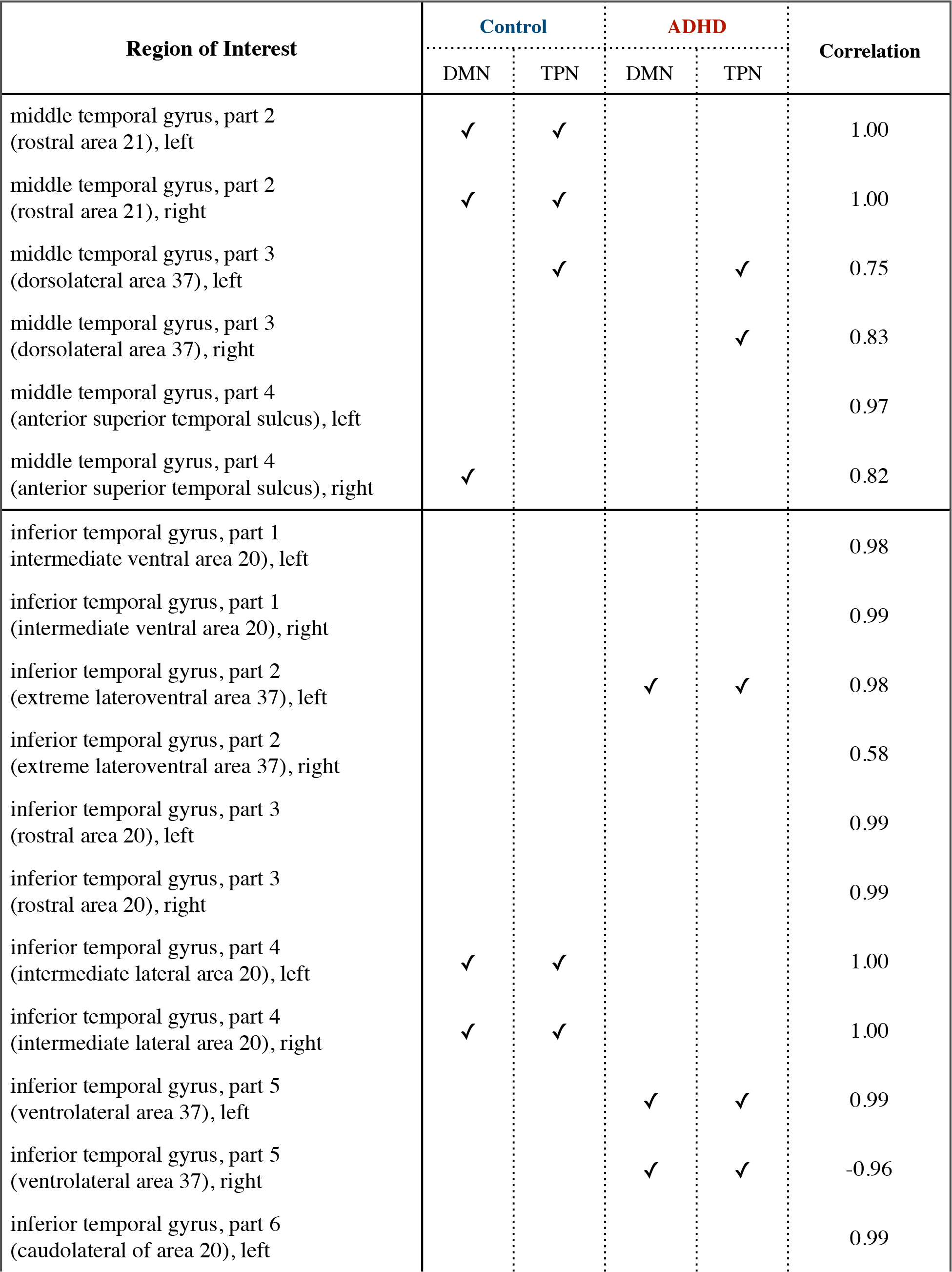

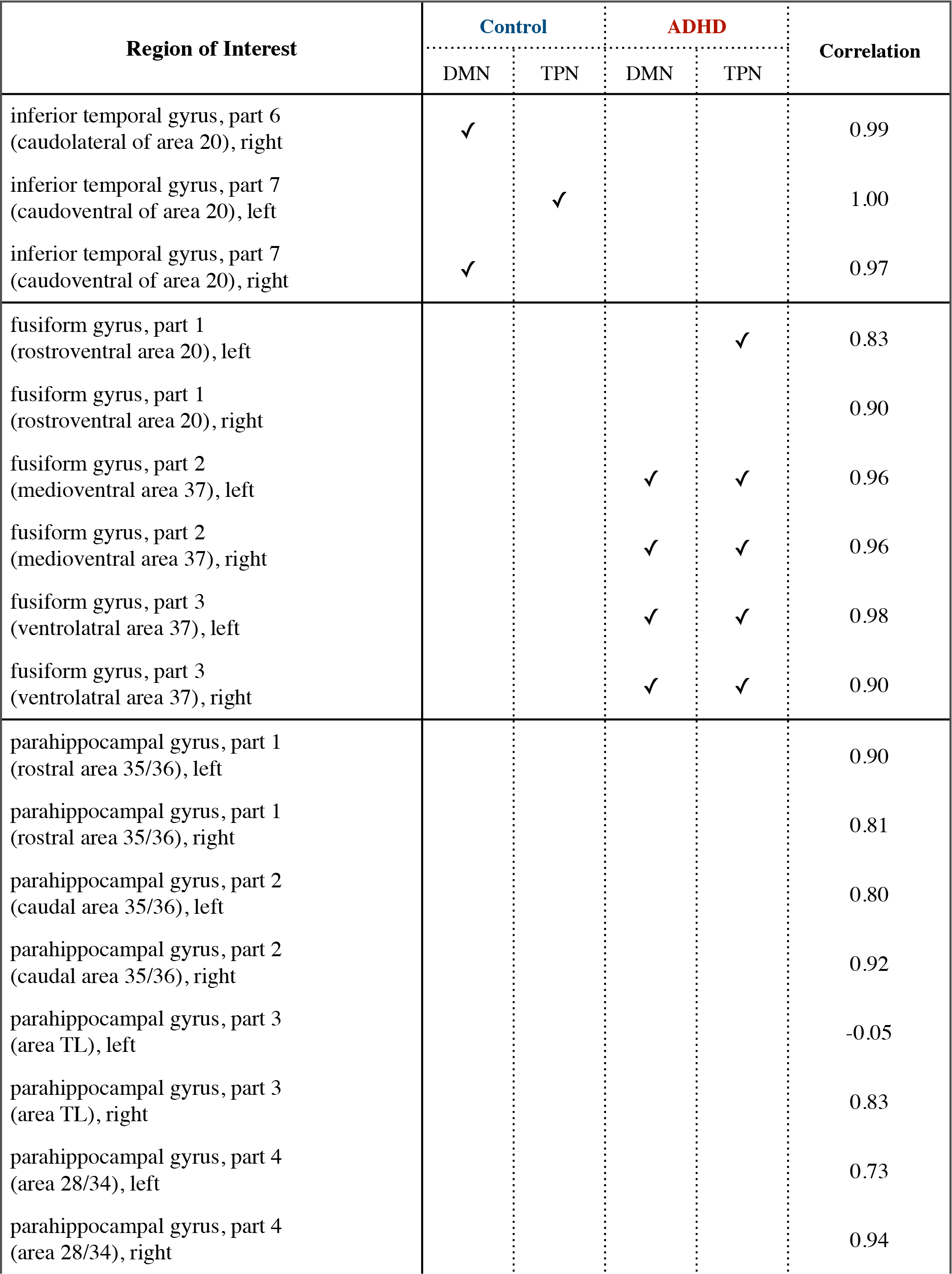

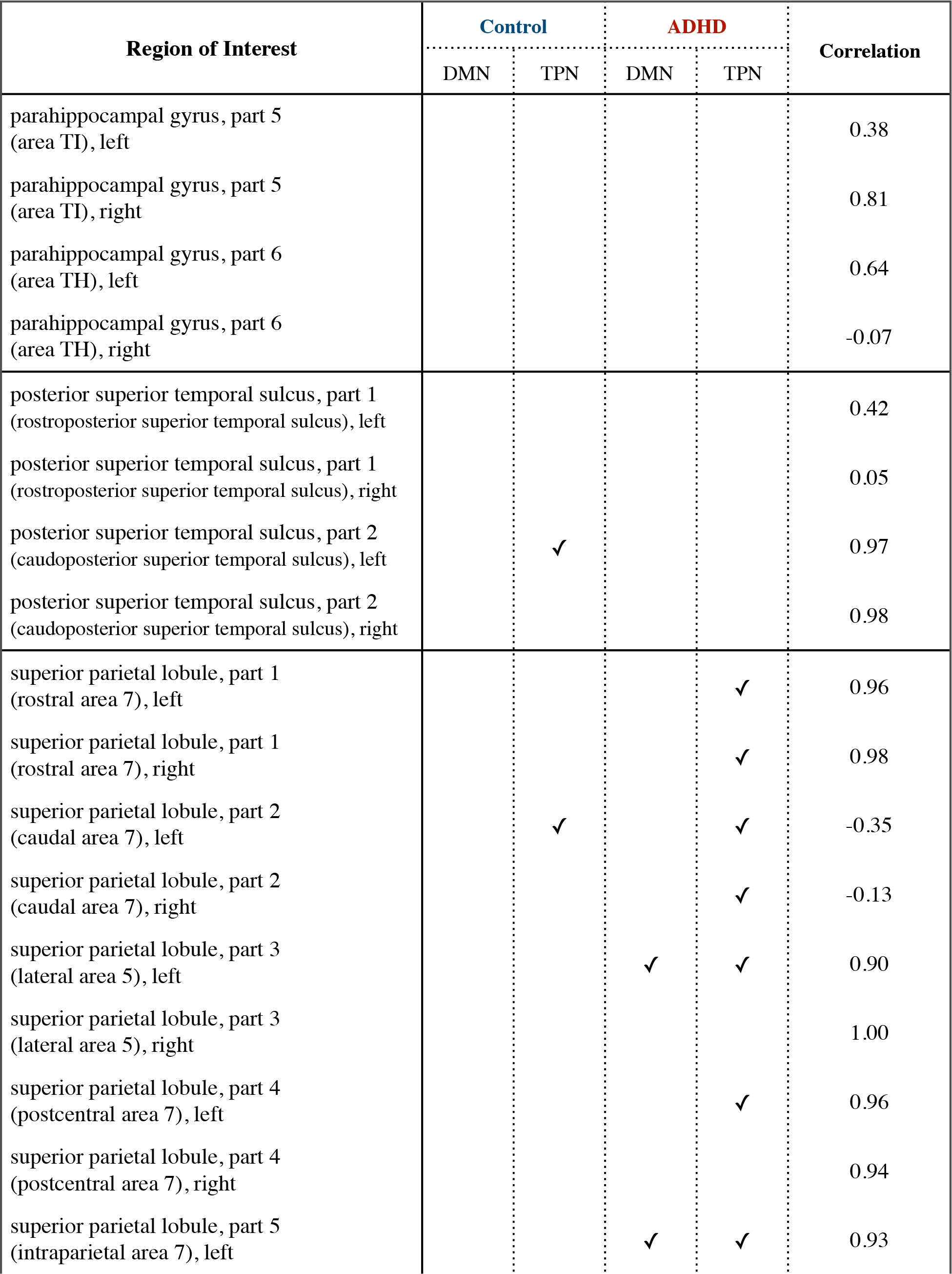

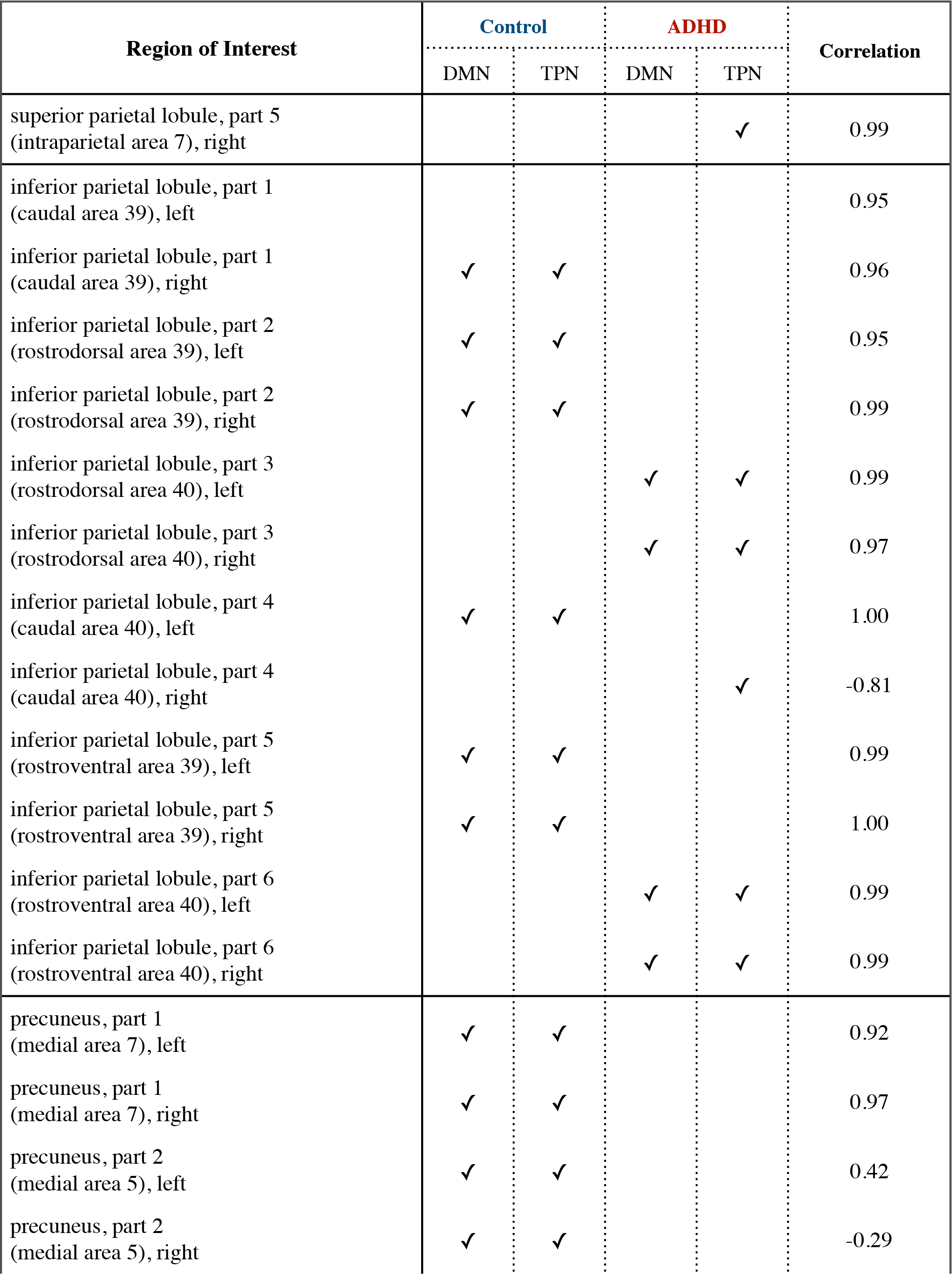

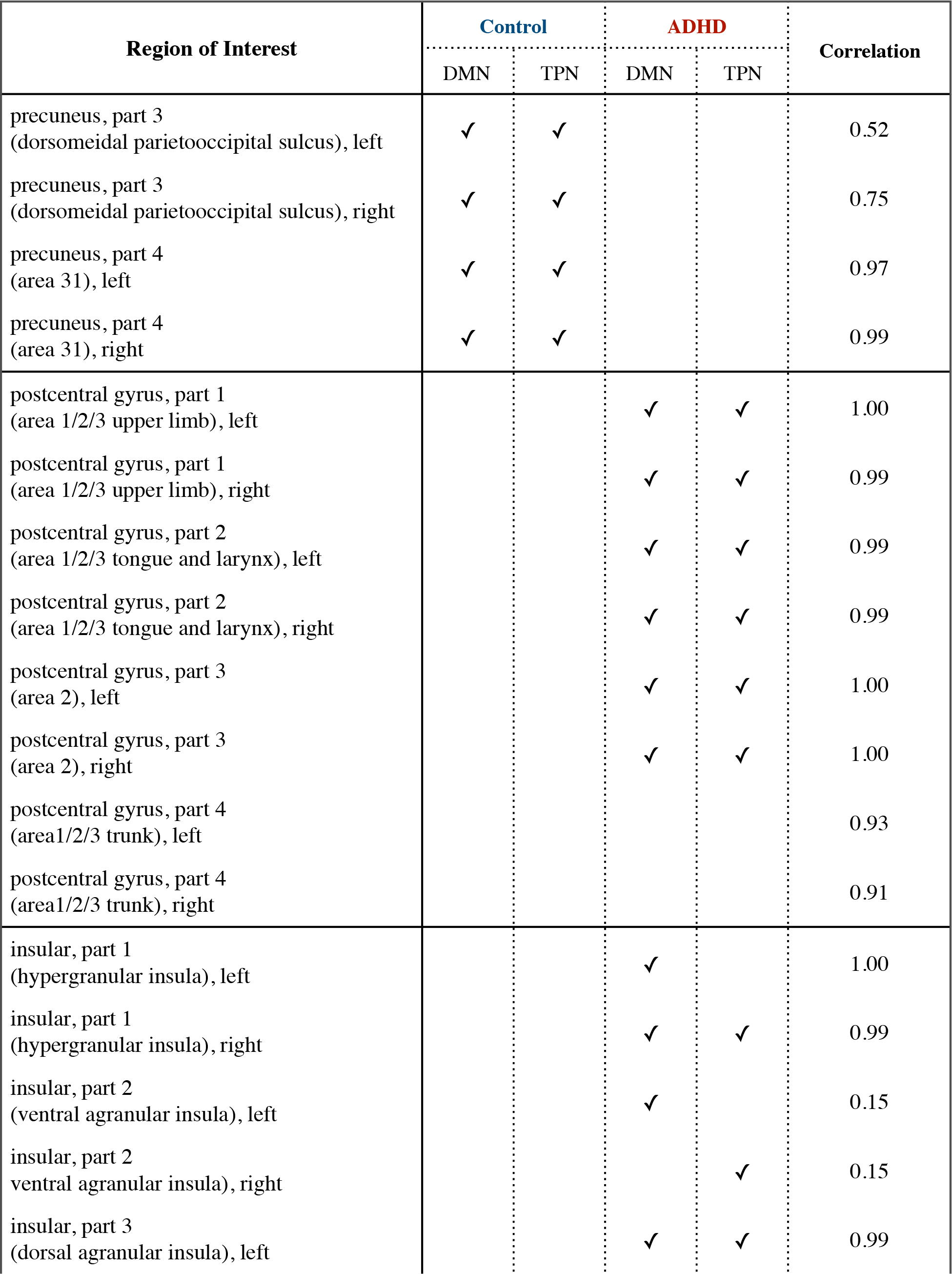

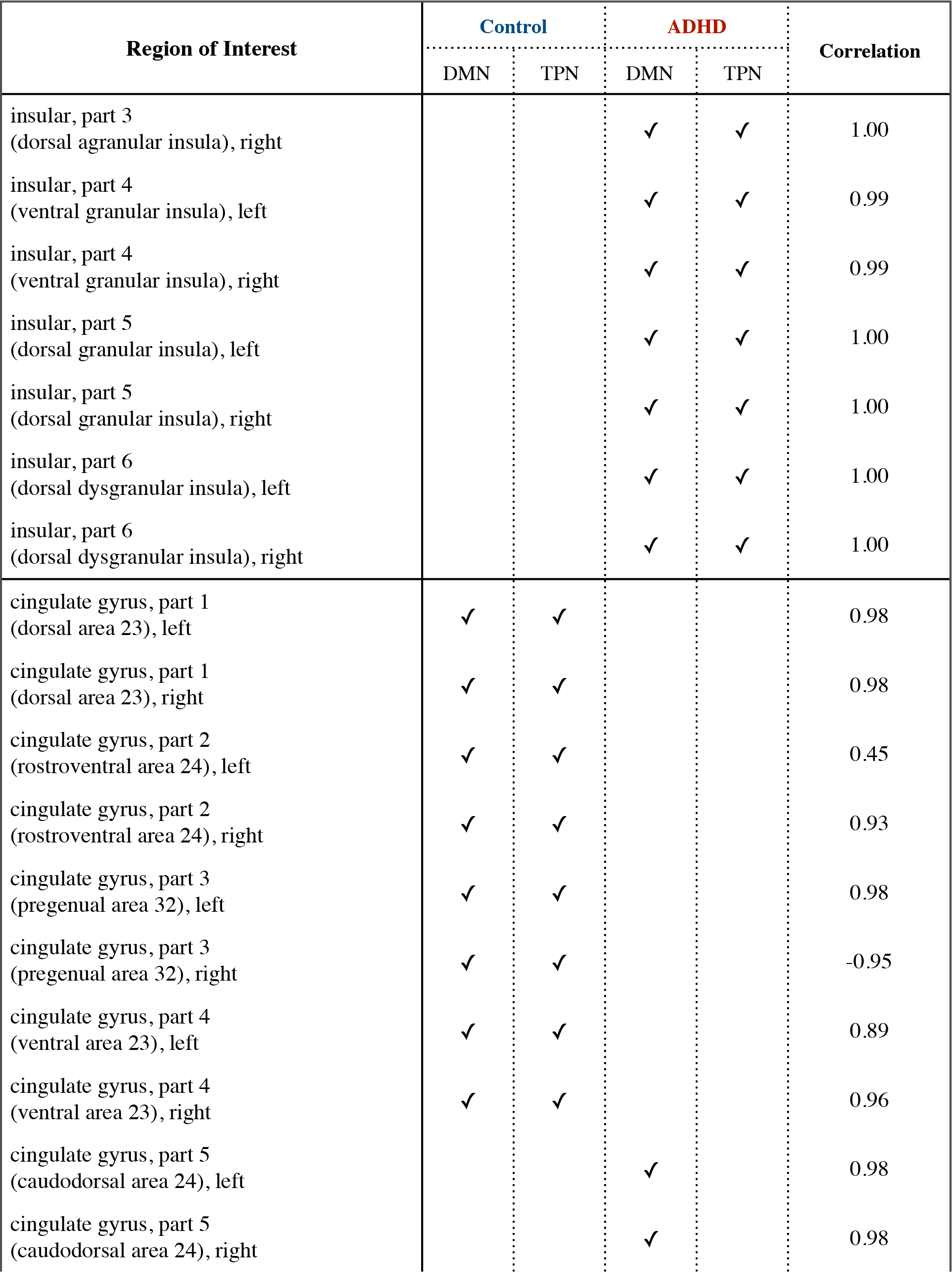

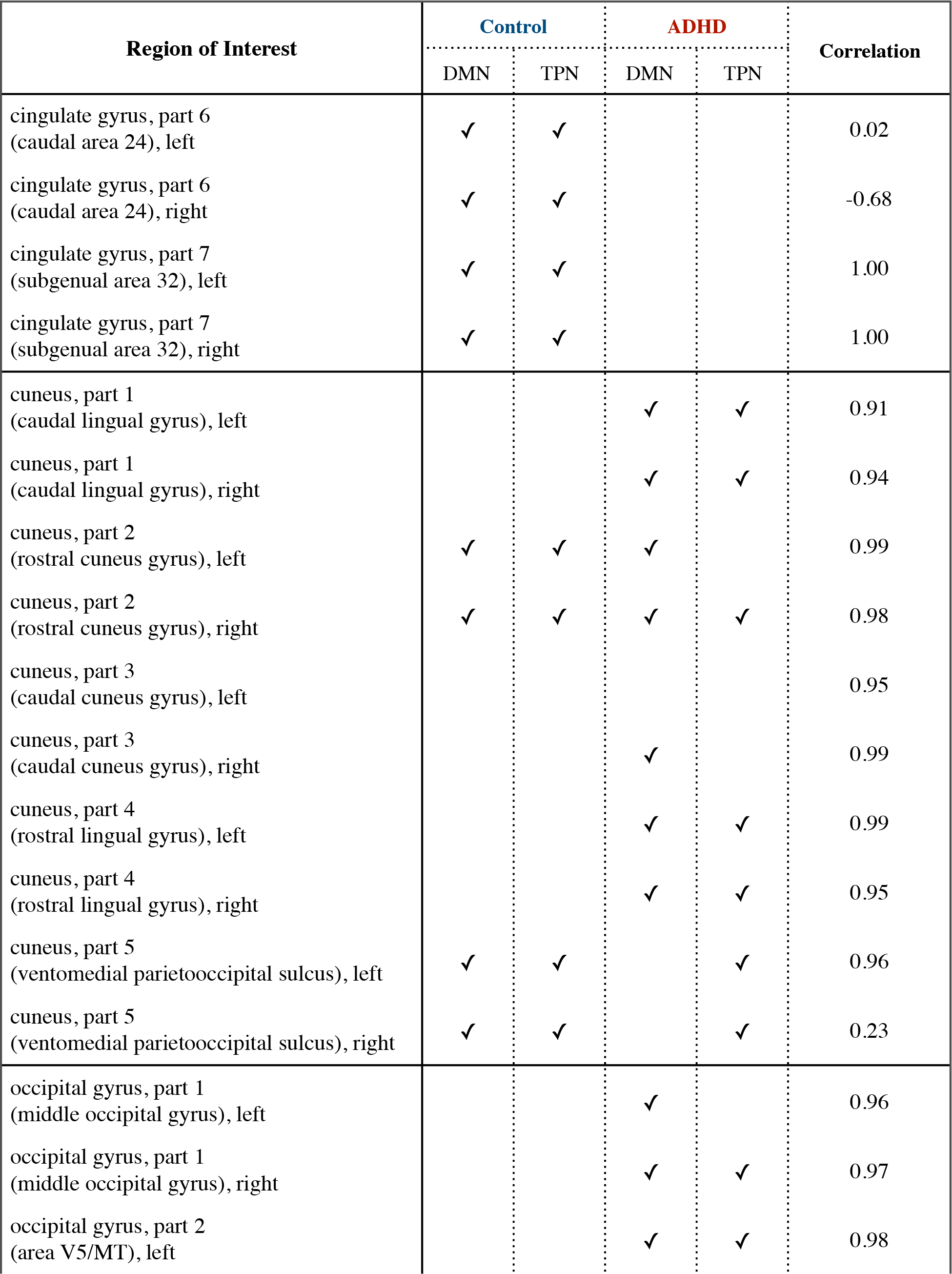

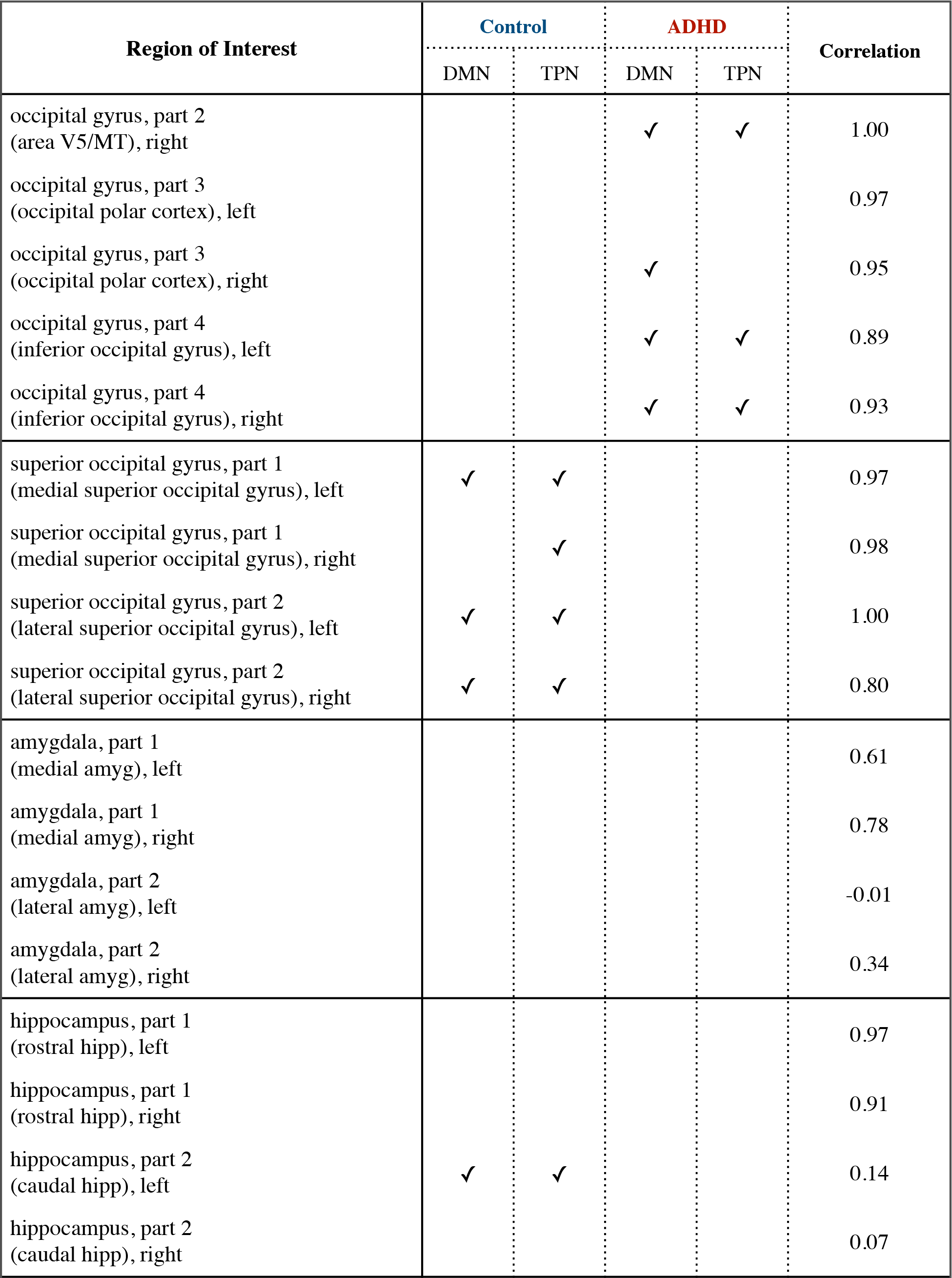

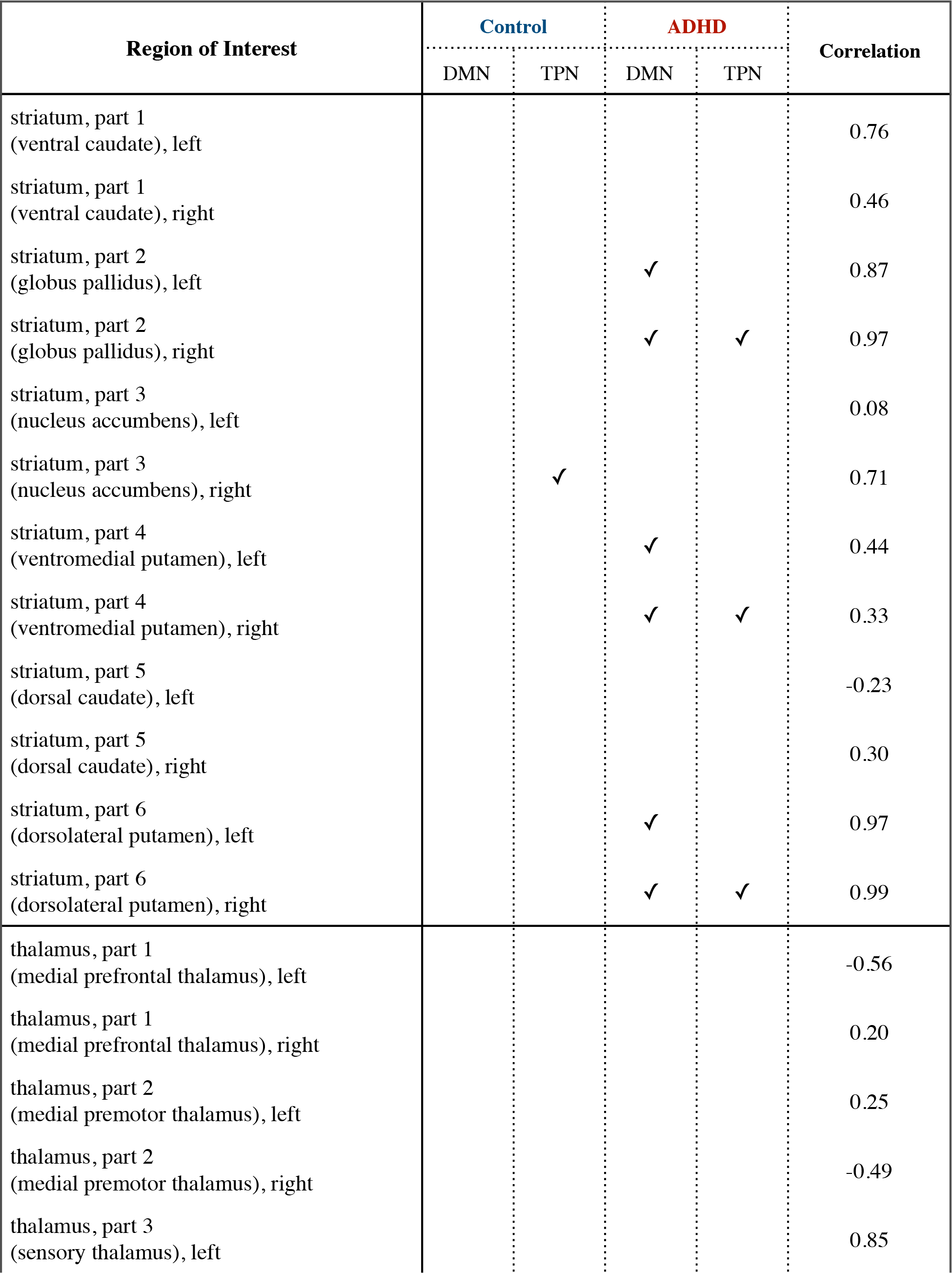

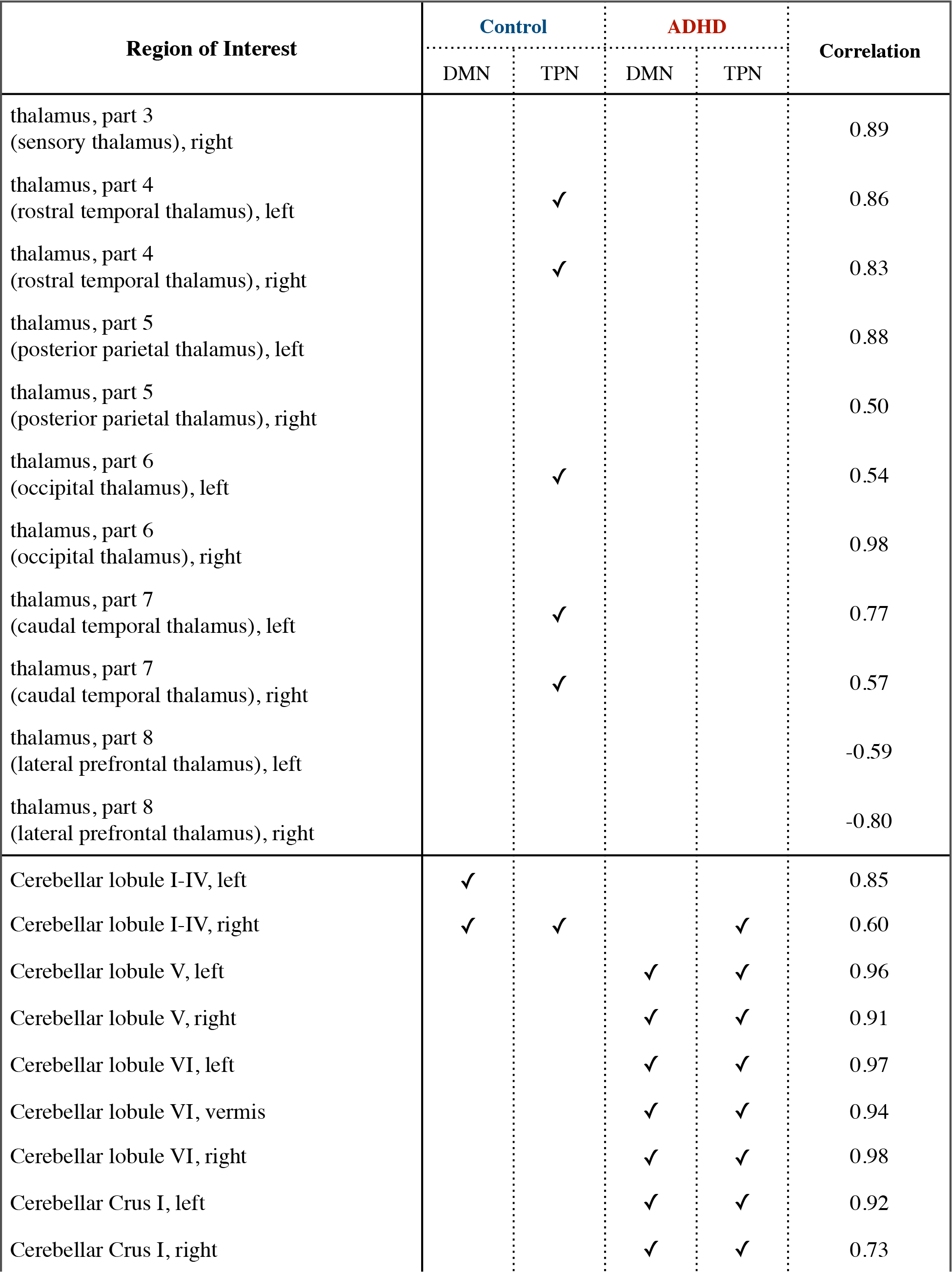

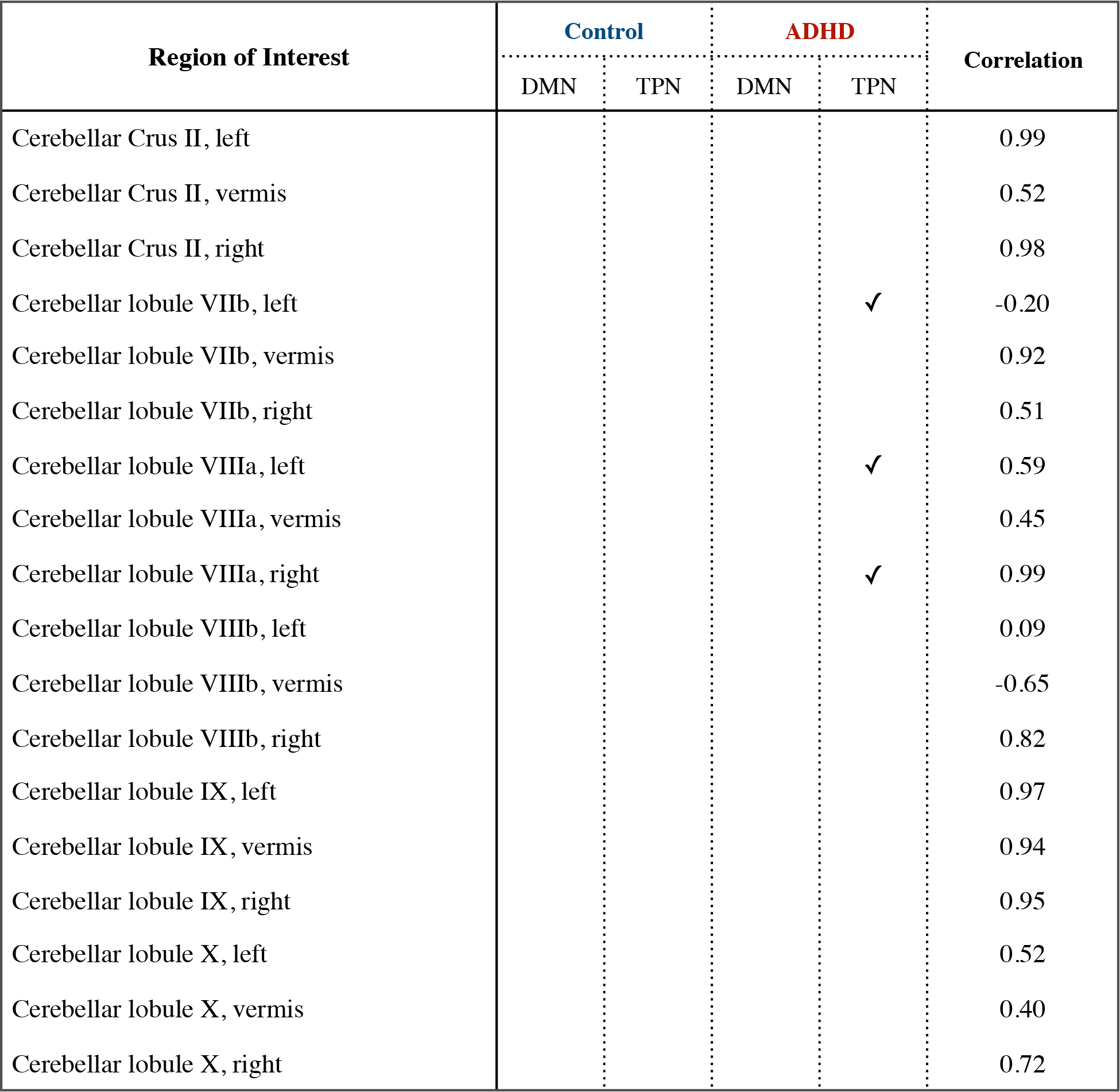

